# Assessing the impact of yield plasticity on hybrid performance in maize

**DOI:** 10.1101/2025.01.21.634104

**Authors:** Jensina M. Davis, Lisa M. Coffey, Jonathan Turkus, Lina López-Corona, Kyle Linders, Chidanand Ullagaddi, Dipak K. Santra, Patrick S. Schnable, James C. Schnable

## Abstract

Improving crop resilience in the face of increasingly extreme and unpredictable weather and reduced access to agricultural inputs such as nitrogen fertilizer and water will require an improved understanding of phenotypic plasticity in crops. To understand the roles of different component traits in determining overall plasticity for grain yield, we generated data from a panel of 122 maize (*Zea mays*) hybrids grown in replicated field trials in 34 environments spanning 700 miles (1126 km) of the U.S. Corn Belt. We observed that the levels of genetic versus environmental control and the relationships between mean parent release year, overall performance, and linear plasticity were trait-dependent across the 18 agronomic and yield components studied. Importantly and unexpectedly, we observed no clear tradeoff between linear plasticity and mean performance and found only rare examples where genotype-by-environment interactions would alter selection decisions based on the environments tested in our dataset. Furthermore, we showed that overall plasticity was repeatable and appears to be under considerable genetic control but that plasticity in response to nitrogen fertilization was not, which may help explain the limited success in breeding for nitrogen use efficiency. Together, these findings improve our understanding of phenotypic plasticity, with implications for maize breeding.

## Introduction

Phenotypic plasticity is the ability of genetically identical individuals to exhibit different phenotypes in response to different environmental conditions. Differences in phenotypic response to the same environments between different genotypes are referred to as genotype-by-environment interactions^1,2^. The proportion of total phenotypic variance that can be attributed to genotype-by-environment interactions varies substantially depending on the population being tested, the environments in which data are collected, and the trait(s) being studied. Phenotypic plasticity can be modeled either as changes in phenotype due to differences in environments, treating the environments as individual factors, or as a relationship between the phenotype and quantitative environmental factors, treating the environments as values along a quantitative axis. In plant breeding, new genetically distinct cultivars have been developed and selected over a period of many years and the final selected varieties will be grown in a wide range of environments over subsequent years. Numerous plant traits of interest to growers exhibit substantial plasticity across environments. The variability in phenotypic plasticity across varieties and traits represents a challenge for plant breeders and farmers because data on how crop varieties perform in one environment will be an imperfect predictor of how these crop varieties will perform in another environment. However, an improved understanding of phenotypic plasticity will help address this challenge by improving predictions.

Changes in climate and agronomic practices also pose challenges for plant breeders and farmers. Historically, plant breeding has occurred in a world of relatively stable environments such that data from numerous locations in past years was able to capture crop performance in a range of environments that were roughly representative of the future environments in which selected crop varieties would be grown. However, as a result of changes in both climate and management practices, the current environments in which new crop varieties are being evaluated are likely less representative of the future environments in which selected crop varieties will be grown. One likely change in management practices is the reduced use of synthetic nitrogen fertilizers due to economic, environmental, and political factors^3–5^. Although it is difficult, if not impossible, to predict and replicate the climactic conditions of the future in our current evaluation environments, it is straightforward to create evaluation environments that consider reduced nitrogen fertilization.

One method of summarizing the phenotypic plasticity of crop varieties is by utilizing model parameters extracted from joint regression models such as Finlay-Wilkinson regression^6^. As discussed above, it is possible to model the environment as either a factor or as a quantitative explanatory variable. In principle, each individual environmental factor likely plays a role in determining phenotypes and different crop varieties may exhibit different responses to different environmental factors. However, given both the extremely large number of potential environmental factors that can vary between experiments and the limited number of total environments in which even the largest replicated field experiments are conducted, it can often be challenging to link variation in specific plant traits to specific environmental factors. Finlay-Wilkinson regression regresses the varietal performance for a given trait across multiple environments on the population-level performance for the trait in a given environment. This jointly estimates the overall genotypic effect as the intercept and its relative response to improved environments as the slope. This method has been widely used to study the genetic architecture of phenotypic plasticity in plants by using the fitted slope as a metric of linear phenotypic plasticity^7–12^.

Breeders may select for different types of yield stability, a concept related to phenotypic plasticity^2^. Type I stability indicates constant performance across environments, and a Type I stable hybrid would be modeled in Finlay-Wilkinson regression as having a linear plasticity value of 0. Type II stable lines have a population mean level of response to improved environments, and would be modeled in Finlay-Wilkinson regression as having a linear plasticity value of 1. Selection for Type I stable hybrids with high yields aims to minimize the effect of genotype-by-environment interactions, but unfortunately, lines with Type I stability are most often low-performing lines. In contrast, selection for Type II stability aims to exploit some of the benefits of genotype-by-environment interactions, while minimizing their deleterious effects. Either of these measures of stability implies selection against hybrids with high linear plasticity values, because of the concern that lines with high linear plasticity values will perform poorly relative to the population in poor environments^1,6^. However, contrasting evidence exists for the relationship between mean trait values and plasticity. In Finlay & Wilkinson (1963)^6^, the authors studied a wheat diversity panel and found a tradeoff between extreme (both high and low) plasticity values and overall performance in terms of yield. Other studies have found either a positive relationship^8,11^ or no relationship^13^ between plasticity and yield, though this may be due to differences in the particular traits studied. For example, Kusmec et al.^7^ found the direction and strength of the relationship between trait mean value and the trait linear plasticity was trait-dependent.

Understanding the contributions of both genotype-by-environment interactions and phenotypic plasticity to crop performance helps to guide decision-making in plant breeding programs today. This understanding will allow us to develop and select cultivars that are most likely to perform well in future environmental conditions. Both phenotypic plasticity and genotype-by-environment interactions have been extensively studied in plant systems^7–11,13–19^. However, the majority of these studies have examined only a few traits^14,15,17,19^, were limited in the range or agronomic relevance of the environments in which varieties were tested^9,11,13,17^, or the used little to no commercially relevant genetic material. Of the studies examining phenotypic plasticity in maize (*Zea mays*), many have focused on identifying genomic regions associated with variation in phenotypic plasticity and most used inbred or doubled-haploid lines^7,9,11,13,15,17^, which are not grown commercially. Studying phenotypic plasticity in a hybrid population allows for an improved understanding of phenotypic plasticity in commercially-relevant germplasm, as most commercial maize production uses hybrid lines. In addition, it is also possible to estimate the contribution of parental inbred lines to the phenotype of the hybrid, termed general combining ability (GCA)^1^.

Part of the complexity of understanding how changes in environment affect crop productivity is that phenotypic outcomes for complex traits such as grain yield are determined by variations in a suite of component phenotypes. Many factors affect grain yield in maize; yield component phenotypes include ears per plant, kernel rows per ear, kernels per row, and mass per kernel. Each of these traits is under partially independent genetic control and each could, in principle exhibit different patterns of response to changes in the environment. Several studies have investigated the phenotypic plasticity of maize yield components^7–11,13^, but only one^10^ used a hybrid population. In addition, how the phenotypic plasticity of maize yield components and agronomic traits differ between older and newer germplasm remains unknown.

In this work, we evaluated the performance of a set of 122 maize hybrids from different breeding eras across up to 34 unique environments representing multiple nitrogen treatments in 10 of the 14 location years for 18 agronomic and yield component traits. Using this dataset, we evaluated the relationships between phenotypic plasticity and performance across environments, with implications for maize breeding programs.

## Results

A set of trait measurements, including combine-measured grain yield, yield components, and a number of other phenotypes were collected from a set of 6,016 four-row maize yield plots. These plots were grown as part of multi-environment field trials of 122 maize hybrids conducted in 2022 and 2023 at field sites in the U.S. Corn Belt near Scottsbluff, Nebraska; North Platte, Nebraska; Lincoln, Nebraska; Missouri Valley, Iowa; Ames, Iowa; and Crawfordsville, Iowa. A number of locations included multiple nitrogen and/or irrigation treatments, resulting in 34 specific environments in which yield and other traits were assayed across the 14 location-years of the study (Figure 1A). Grain yields (14.55 – 238.12 bushels/acre (0.91 – 14.95 MT/ha), Figure 1B) were representative of the range of yields within 95% confidence intervals of county mean yields across Nebraska and Iowa in 2022 and 2023 (52.02 – 254.15 bushels/acre (3.27 – 15.96 MT/ha)^20^). Of the 122 hybrids, 81 were evaluated in all 34 envronments. A total of 18 unique phenotypes were scored in these experiments. After the removal of extreme values (0.17 – 2.97% per phenotype), the resulting dataset contained 94,053 non-missing phenotype values.

**Figure 1.**
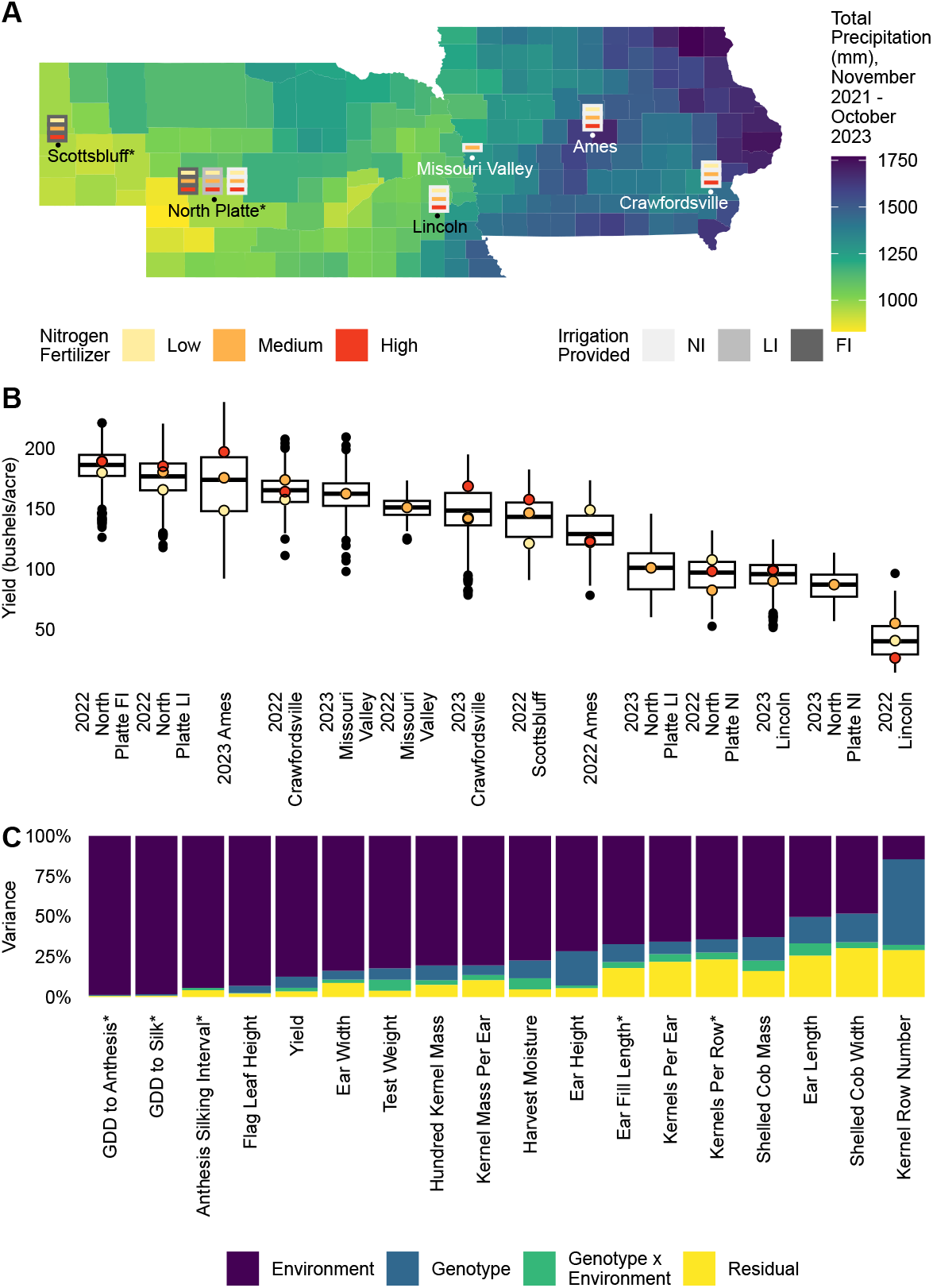
Multi-environment hybrid maize field trials across Nebraska and Iowa show a large effect of environment on yield component traits aside from kernel row number. **A)** An overview of the experimental design utilized in this study overlaid on a map of Nebraska and Iowa, showing the total precipitation during the period relevant to the growing seasons in this study (2022 – 2023) from the approximate end of the 2021 growing season (November 1, 2021) to the approximate end of the 2023 growing season (October 31, 2023). The experimental design included three nitrogen fertilization treatments (Low: 75 lb N/acre (84 kg N/ha), Medium: 150 – 175 lb N/acre (168 – 196 kg N/ha), and High: 225 – 250 lb N/acre (252 – 280 kg N/ha)) and three irrigation treatments (No Irrigation (NI): 0 mm, Limited Irrigation (LI): 100 – 200 mm, and Full Irrigation (FI):*>* 200 mm) for the two study years. *The Scottsbluff site was used in the 2022 growing season and, in North Platte, the experimental design shown applies only to the 2022 growing season. In 2023, a single field was grown in North Platte with two irrigation treatments (NI = 0 inches (0 mm) and LI = 4.5 inches (114 mm)) in a randomized incomplete block design with 150 lb N/acre (252 kg N/ha) nitrogen fertilizer applied. **B)** Population-level yield in each location-year (*n* = 135 – 560 plots). The colored points indicate the median yield of the corresponding nitrogen treatment within the location-year. Box plots indicate the range from the 25^th^ – 75^th^ percentile of values. Black lines within the boxplots indicate the median value. Whiskers indicate the most extreme values within 1.5 times the interquartile range and black points indicate the values of data points outside that range. **C)** Estimated proportion of variance explained by environment, genotype, and genotype-by-environment interactions for each of the 18 phenotypes measured. Asterisks denote phenotypes measured only in a subset of location-years. GDD indicates growing degree days.

Differences between environments comprised the largest contributing factor to the variation of 17 of the 18 phenotypes based on a variance partitioning analysis including environmental, genotypic, and genotype-by-environment interaction factors (Figure 1C). Only for kernel row number were differences between genotypes the most significant contributor to variation (53.2%). Differences between environments explained more than 98% of the variation in growing degree days (GDD) before anthesis and in GDD to silking and they explained more than 70% of variation in flag leaf and ear heights. Genotype-by-environment interactions did not explain the largest proportion of variation in any of the phenotypes tested, but the phenotypes for which this factor played the largest role were ear length (7.7%), test weight (6.9%), and harvest moisture (6.9%). Although numerous components contribute to the environment as a factor as in the variance partitioning model used here, one well-known component is nitrogen fertilization, which is expected to have a positive relationship with grain yield.

In three of the ten location-years with nitrogen treatments (2023 Ames, 2022 Scottsbluff, and 2022 North Platte LI), we observed a positive relationship between nitrogen fertilization rate and yield at the population level (Figure 1B). We fit the Finlay-Wilkinson model for the three environments contained within each of these three location-years to determine the repeatability of nitrogen responses across location-years. Linear plasticity for yield across nitrogen application rates did not correlate across the location-years studied, with Spearman rank correlations for location-year pairs ranging from -0.07 to 0.15 (Figure 2A). Similar trends were observed for hundred kernel mass (Figure 2B) and other traits (Supplemental Figure 1).

**Figure 2.**
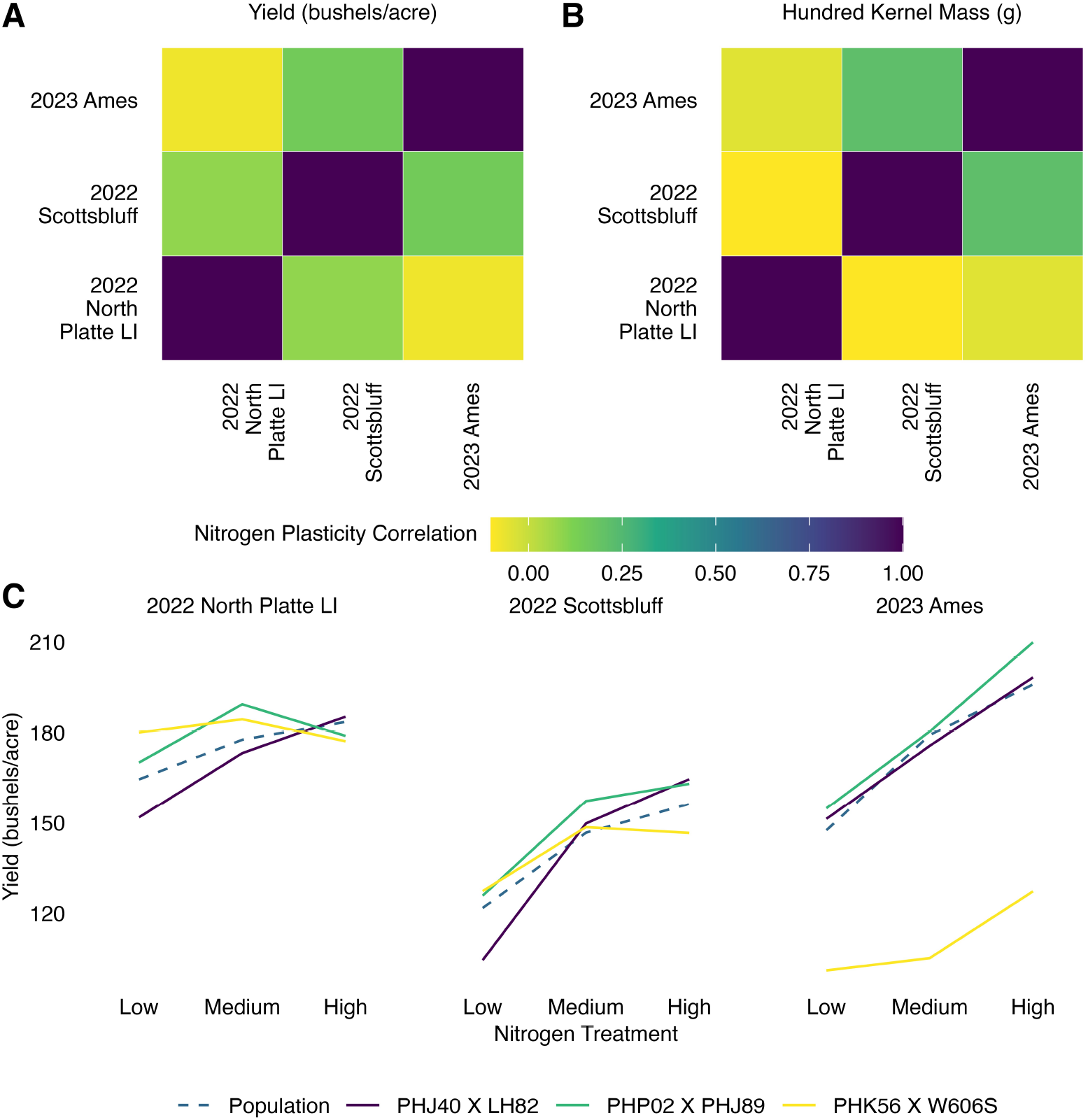
The responses of maize hybrids to nitrogen fertilization differ between location-years. **A)** Spearman rank correlations of yield linear nitrogen plasticity values across all hybrids common to both location-years for location-years where a population-level positive relationship of yield with nitrogen fertilization was observed. **B)** Spearman rank correlations of hundred kernel mass linear nitrogen plasticity values across all hybrids common to both location-years for location-years where a population-level positive relationship of yield with nitrogen fertilization was observed. **C)** Mean yields of hybrids with the lowest mean nitrogen plasticity (MNP) value (PHK56 X W606S, MNP = 0.89), an intermediate MNP value (PHP02 X PHJ89, MNP = 1.00), and the highest MNP value (PHJ40 X LH82, MNP = 1.11) and population-level response under different nitrogen fertilization treatments in location-years where a population-level positive relationship of yield with nitrogen fertilization was observed.

The nitrogen responses of individual lines to increased nitrogen fertilization differed between location-years (Figure 2A, B), and sometimes within a location-year (Supplemental Figure 2). We selected three lines having the lowest mean nitrogen plasticity (MNP) value (PHK56 X W606S, MNP = 0.89), an intermediate MNP value (PHP02 X PHJ89, MNP = 1.00), and the highest MNP value (PHJ40 X LH82, MNP = 1.11) across the three location-years (Figure 2C). All three hybrids exhibited multiple patterns of nitrogen response across the three location-years (Figure 2C), including differential responses to increasing nitrogen fertilization from low to medium levels and from medium to high levels. The repeatability of nitrogen plasticity values within a location-year for yield was quite low (Spearman *ρ* = 0.0473, Supplemental Figure 2) when comparing nitrogen plasticities estimated with one block per nitrogen treatment in each of the three location-years where the population-level response to nitrogen was positive. For the maize hybrids in our population, both within and across location-years, the relative nitrogen plasticity for yield showed poor repeatability (− 0.07 ≤ *ρ* ≤ 0.17), and it often differed from the population-level nitrogen response in the given location-year (Figure 2C).

Given our limited success in studying the response of maize hybrids to a specific environmental factor, we used Finlay-Wilkinson joint regression to investigate the relative response to improved environments of the maize hybrid lines in our population, without regard to the specific environmental factor(s) that improved the environment. We chose to exclude the 2022 Lincoln location-year from Finlay-Wilkinson regression analyses given its extremely low yield value relative to all other locations (Figure 1B). We found a moderate correspondence (Spearman *ρ* = 0.4483, Supplemental Figure 3) between the linear plasticity values for each hybrid estimated by two Finlay-Wilkinson models each using half of the dataset containing one block from each environment. This correspondence is within the range of the Spearman correlations between blocks for the 18 traits directly measured in this study (0.4411 – 0.9559). This indicates that the linear plasticity values extracted from Finlay-Wilkinson regression models across all 31 environments used are reasonably repeatable and represent a genetically controlled trait.

Increased phenotypic plasticity is typically assumed to be undesirable, because of the concern that lines with high plasticity will perform particularly poorly in poor environments. Figure 3A shows the Finlay-Wilkinson yield regression line for the hybrids with the highest, intermediate, and lowest Finlay-Wilkinson linear plasticity values (slopes) across the tested environments. Of these three lines, the one with the highest plasticity is predicted to have the highest yield in all tested environments, followed by the line with intermediate plasticity, indicating that increased linear plasticity may not be associated with poor performance in poor environments.

**Figure 3.**
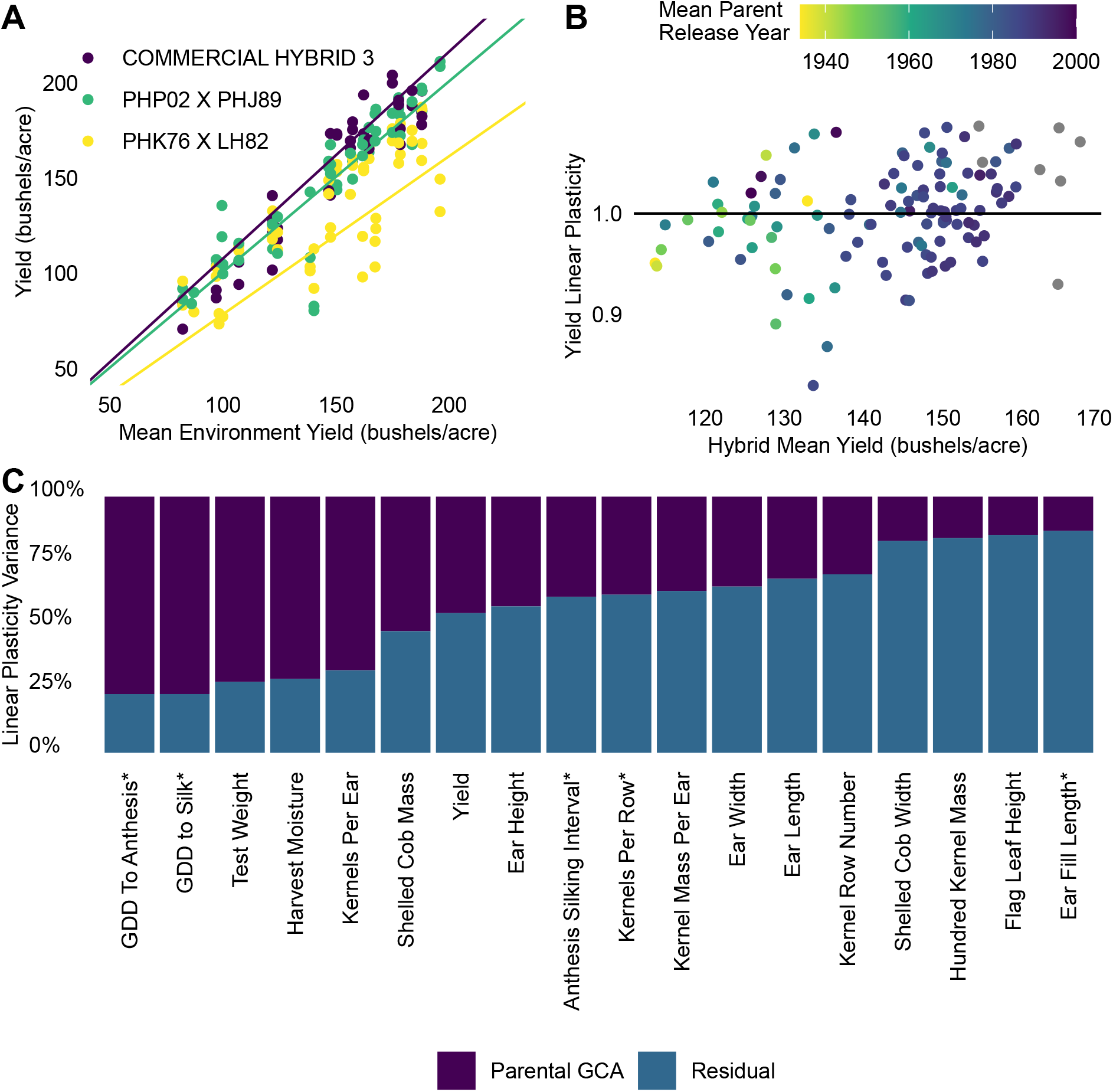
Increased linear plasticity is associated with higher performance across all environments. **A)** Performance of lines with the highest (COMMERCIAL HYBRID 3, *b* = 1.09), an intermediate (PHP02 X PHJ89, *b* = 1.00), and the lowest (PHK76 X LH82, *b* = 0.83) Finlay-Wilkinson plasticity values for yield versus mean yield of each environment in this study and their corresponding Finlay-Wilkinson regression lines. **B)** Finlay-Wilkinson yield linear plasticity values for each hybrid in this study versus their mean yields across all environments, colored by their mean parent release year. Gray points represent commercial check hybrids for which the inbred parents are unknown. The black horizontal line indicates a linear plasticity value of 1. **C)** Proportion of the linear plasticity of yield component traits attributable to the general combining ability (GCA) of the hybrid parents. Parental GCA represents the sum of variances attributed to either the ear parent or the pollen parent GCA. Asterisks denote phenotypes measured in only a subset of location-years. GDD indicates growing degree days.

In our dataset, yield and linear plasticity were significantly positively correlated (*p* = 1.1 × 10^−3^). Hybrids with later mean parent release years tended to have higher yields (*p* < 2.2 × 10^−16^), indicating that the association between grain yield and linear plasticity may be due to the effect of artificial selection. The significance of the association between yield and linear plasticity increases marginally when controlling for mean parent release year (*p* < 3.6 × 10^−4^, Figure 3B), which ranged from 1934 to 2000 in our dataset. However, the performance of the most plastic or the highest yielding hybrids as a percentage of the environment mean did not drastically increase as the environment mean increased (Supplemental Figure 4), as would be expected if the increases in linear plasticities were independent of the increases in performance across all environments. The increases in linear plasticity and their positive relationship with overall yield and mean parent release year are at least partially due to increased performance across all environments (Supplemental Figures 5, 6), rather than being solely due to an increase in the level of response to favorable environments.

In contrast to the pattern observed for grain yield, the kernel row number, ear length, ear fill length, kernels per row, and kernels per ear did not show significant associations (*p >* 0.05) between mean value and either linear plasticity or mean release year of inbred parents (Supplemental Figure 7. Mean trait values for ear width (*p* = 0.01), hundred kernel mass (*p* = 0.04), kernel mass per ear (*p* = 3.7 × 10^−5^), and harvest moisture (*p* < 2.0 × 10^−16^) were all significantly positively correlated with linear plasticity, but none of these traits exhibited a significant correlation between mean value and mean release year of the inbred parents (Supplemental Figure 7). Flag leaf height, shelled cob width, and shelled cob mass had no significant association with linear plasticity (*p >* 0.05), but they had significant negative associations with mean parent release year (*p* < 1.0 × 10^−3^), indicating that they have declined over time. For test weight, the trait mean has significantly increased over time (*p* = 0.05), but lower linear plasticity was significantly associated with higher overall mean trait values (*p* < 2.0 × 10^−16^), indicating potential indirect selection for high and stable test weights. The overall trait means of ear height (*p* = 4.2 × 10^−15^), GDD to anthesis (4.5 × 10^−6^), GDD to silk (*p* = 1.9 × 10^−7^), and anthesis-silking interval (*p* = 0.02) have all significantly declined over time. Increased linear plasticity was associated with decreased overall mean anthesis-silking intervals (*p* = 3.7 × 10^−7^), whereas decreased linear plasticities for ear height (*p* = 3.6 × 10^−3^), GDD to anthesis (*p* = 2.0 × 10^−4^), and GDD to silk (*p* = 3.2 × 10^−3^) were associated with decreased overall mean trait values. The mixed relationships between how the traits have changed over time and linear plasticity with overall trait values indicate that increased or decreased linear plasticities are not universally neutral, advantageous, or disadvantageous for overall performance, but rather, are trait-dependent (Supplemental Figure 7).

Having observed the significant relationship between mean parent release year and linear plasticity, we investigated how much variation in linear plasticity could be explained by the particular parental lines used. Parental plasticity general combining ability (GCA) explained the most variation in the linear plasticity of GDD to anthesis (77.1%), followed by GDD to silk (77.1%) and test weight (72.3%, Figure 3C), and contributed the least to the variance in the linear plasticity of flag leaf height (14.9%) and ear fill length (13.3%). We found that 45.5% of the variation in the yield linear plasticity could be explained by variation in parental GCA. The relatively large contribution of parental GCA to the linear plasticity of GDD to anthesis, GDD to silk, and test weight indicates the promising potential to modify the level of linear plasticity of hybrids for these traits via parental selection though it may be more difficult to modify the level of linear plasticity for flag leaf height and ear fill length.

Genotype-by-environment interactions that result in a change in rank ordering between hybrids within the population of environments have a greater effect on selection decisions within breeding programs than do genotype-by-environment interactions that do not result in rank order changes. We found that 14.6% of hybrid pairs were predicted to change rank order between the environments with the lowest and highest population mean yields in the 31 environments used to fit the Finlay-Wilkinson model using their regression-fitted values (Figure 4A), and that 32.6% of predicted rank order changes were for hybrid pairs with mean yields separated by more than 10 bushels per acre (0.63 MT/ha) in the most favorable environment. The maximum distance between overall ranks for hybrid pairs predicted to have a rank order change was 48 (Supplemental Figure 8). The hybrids in this pair were predicted to have a mean yield difference of 29.0 bushels per acre (1.82 MT/ha) in the most favorable environment by Finlay-Wilkinson regression. However, their observed mean yields in the most favorable environment differed by 56.9 bushels per acre (3.57 MT/ha) and their overall mean yields across the 31 environments differed by 16.9 bushels per acre (1.06 MT/ha). Although this analysis gives useful insight into how often rank order changes may occur because of genotype-by-environment interactions within this population of environments, it does not indicate how often they represent meaningful differences between hybrids that may result in different selection decisions depending on the environments used for selection.

**Figure 4.**
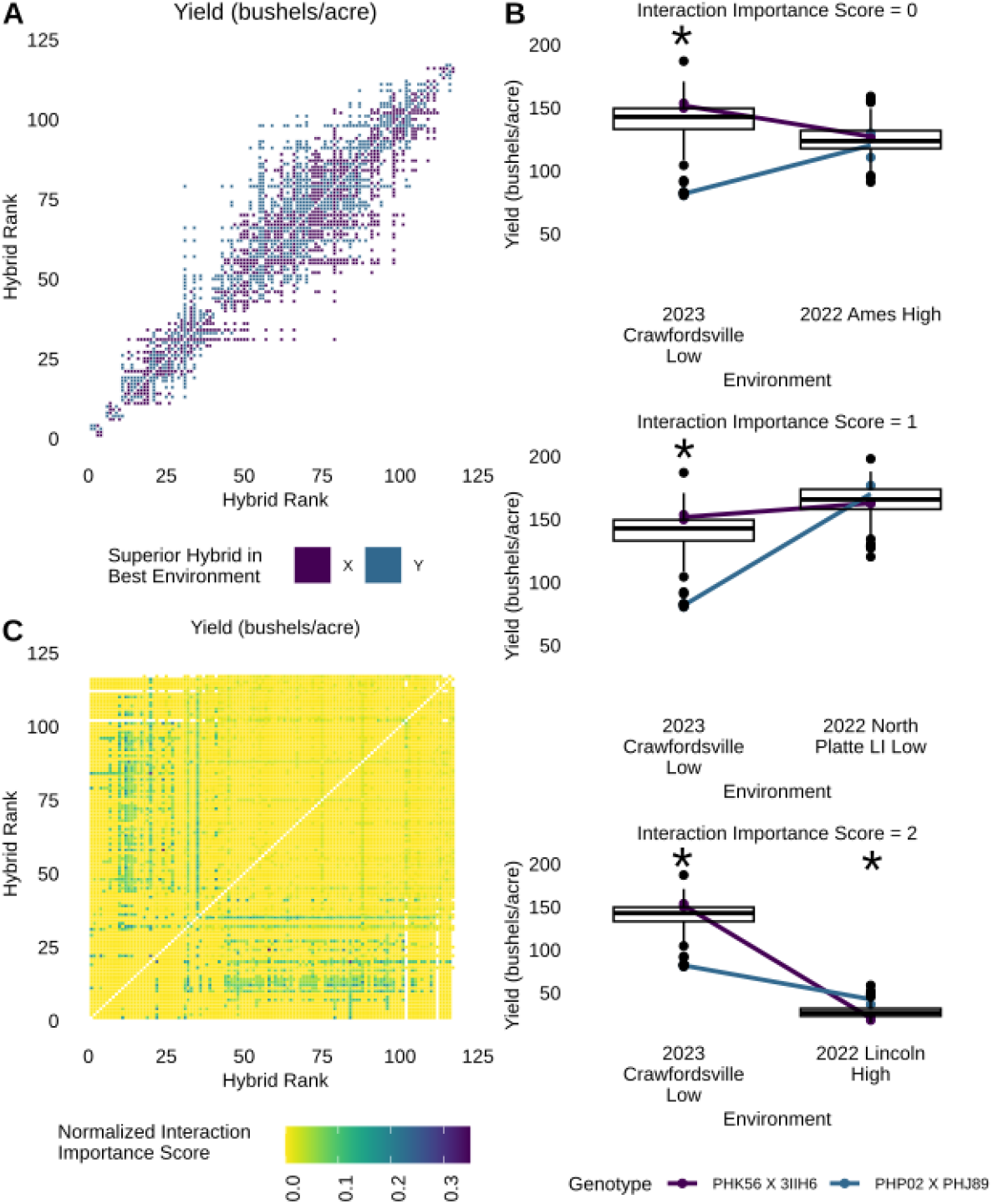
Importance of genotype-by-environment interactions to selection decisions regarding a pair of hybrids depends on the environments used for selection. **A)** Incidence matrix indicating whether Finlay-Wilkinson regression predicts that the two hybrids will change rank order between the lowest and highest population mean environments used to fit the model. Hybrids are ranked in order of ascending yield BLUP values fitting the environment as a fixed effect. The colored squares indicate a rank order change was predicted for the hybrid pair and the color indicates which hybrid is predicted to have a higher yield in the environment with the highest population mean yield. **B)** Performance and genotype-by-environment interactions of one hybrid pair across three environment pairs. Box plots show population-level yield in the given environment (*n* = 163 – 175 plots). Boxes indicate the range from the 25^th^ – 75^th^ percentile of values. Black lines within the boxplots indicate the median value. Whiskers indicate the most extreme values within 1.5 times the interquartile range and black points indicate the values of data points outside that range. The colored lines connect the mean yields of each hybrid between environments. Asterisks denote a significant difference in mean yields between the two hybrids in the environment at a significance level of *α* = 0.05 according to Tukey’s HSD. The top panel shows a pair of environments where the hybrid pair did not change rank order and received an interaction importance score of 0. In the middle panel, the hybrids changed rank order in this pair of environments and had a significant difference in mean yield in one of the two environments, and they received an interaction importance score of 1. The bottom panel shows an environment pair where the hybrids changed rank order and had significant differences in mean yield in both environments, and they received an interaction importance score of 2. **C)** Incidence matrix indicating the frequency with which two hybrids exhibited an interaction for yield between two environments that represents a potentially important change in the selection decision between environments. For each hybrid pair, interaction scores were summed across all environment pairs and divided by the total score possible for the hybrid pair based on the number of environments in which both hybrids were present. Hybrids are ranked in order of ascending yield BLUP values fitting the environment as a fixed effect.

The hybrids 2369 X LH123HT (overall rank 58, in order of ascending BLUP) and K201 X OS426 (overall rank 24) exhibited rank order changes in four of the six environment pairs where both hybrids were present, with a significant difference in one of the two environments for pairs of environments, all of which had medium nitrogen application rates (2023 Ames and 2023 Missouri Valley; 2023 Ames and 2023 North Platte NI; 2023 Crawfordsville and 2023 Missouri Valley; 2023 Crawfordsville and 2023 North Platte NI). The change in rank order combined with the significant difference in one environment produced an interaction importance score of 1 for each of these four environment pairs (Figure 4B), and a total interaction importance score equal to 33.3% of the total possible score for this hybrid pair. We estimated the interaction importance scores for all hybrid pairs in all environment pairs where both hybrids were present for all 18 traits, and we provide a detailed description of this method in Methods. Eight other hybrid pairs also had normalized interaction importance scores ≥ 0.20, but these were extremely rare cases representing only 0.13% of all hybrid pairs. Out of 6,786 hybrid pairs, 17.0% of hybrid pairs exhibited no important interactions, with either no rank order changes across the tested environments or no significant difference in mean yields when a rank order change did occur. Furthermore, 92.5% of hybrid pairs had interaction importance scores less than 5% of the maximum score (Figure 4C), indicating that few of the genotype-by-environment interactions would result in a significantly different selection decision had a different environment been used for selection. However, the pattern and level of interaction importance scores varied substantially among traits (Supplemental Figure 9), with some traits such as GDD to silk having interaction importance scores as high as half of the total possible score, but being quite rare and concentrated in hybrid pairs where both hybrids were ranked in the upper 50% of hybrids for the trait (Supplemental Figure 9D). Normalized interaction importance scores ≥ 10% of the total possible score were much more widespread for test weight and harvest moisture than for any other traits, with 9.2% and 7.9% of hybrid pairs in our population, respectively, having normalized interaction importance scores ≥ 0.10 for these traits (Supplemental Figure 9F, G). Normalized interaction importance scores were lower for kernel mass per ear than for yield (Supplemental Figure 9Q), indicating both that important genotype-by-environment interactions for kernel mass per ear are rarer than for yield, and that plasticity in ear phenotypes may not fully explain plasticity in overall grain yield. It is also notable that few of the hybrid pairs with a relatively high interaction importance score for yield overlapped with the hybrid pairs predicted by Finlay-Wilkinson regression to have a rank order change for yield in the set of environments tested (Figure 4A, C). This may be due to similar performance between many pairs of hybrids that were predicted by Finlay-Wilkinson to have rank order changes (Figure 4A, Supplemental Figure 8), indicating that few of the Finlay-Wilkinson predicted rank order changes between hybrids in our environments would have actually resulted in different selection decisions.

## Discussion

Breeders may take several approaches toward genotype-by-environment interactions– ignoring them or, most often, minimizing them or exploiting them. Approaches to genotype-by-environment interactions in breeding programs that seek to minimize or exploit them assume that they will have important impacts on selection decisions. Hybrid pairs in our population rarely had interactions that would significantly change selection decisions due to rank order changes with significant differences in yield in one or more of the environments under comparison (Figure 4C). Though some hybrids in the lower 50% of the population for yield had important genotype-by-environment interactions with a majority of hybrids in the upper 50% of the population for yield, these hybrids represented only 14.5% of our population. The low frequency of genotype-by-environment interactions that would change selection decisions revealed an opportunity to reduce the number of environments used for selection. However, it is possible that these results would not translate to an active breeding population. Our population of hybrids was generated using inbred parental lines that were historically patented by either public or private breeding programs over a 68-year period with hybrid mean parent release years ranging from 1934 to 2000. Because hybrid mean yield has increased over this time period (Figure 3B, Duvick et al.^21^), our population may show larger differences in mean performance between genotypes than would a population of elite hybrids originating from the same selection cycle in an active breeding program. This may lead us to observe fewer genotype-by-environment interactions that would likely directly affect selection decisions.

The differences between our observation that hybrid mean yield and linear plasticity are positively associated and observations from previous studies indicating a tradeoff between them may be due to several factors. First, it may be due to differences in the populations sampled, especially for studies, such as ours, that utilized stability measures derived from joint regression models that are indexed on the population used to fit the model (Figure 3A). Our population of hybrids was generated using inbred parental lines historically patented by either public or private breeding programs; thus these parental lines represent some of the most elite parental lines existing in breeding programs at the time of patent issuance. Second, our population includes hybrids with mean parent release years ranging from 1934 to 2000, so some effects observed here may be artifacts of different time periods of crop improvement. Third, many studies^6–8,11^, including this study, have indexed the regression based on the range of mean trait values in the range of environments studied, which may also contribute to the low agreement among studies. However, it is also possible that, to some extent, the association we observed between high average yield and high plasticity is an artifact of the way plasticity is calculated. Yield as a percentage of the environment mean of either the most plastic or highest overall yielding hybrids does not dramatically increase as the environment mean increases (Supplemental Figure 4), indicating that the increase in linear plasticity may not be independent of increased performance across all environments.

When we estimated linear plasticity for yield as a percentage of the environment mean, we observed a tradeoff between linear plasticity and both average performance relative to the population and average yield across all environments (Supplemental Figure 10A, B), as was initially expected. It is typically assumed that a tradeoff between plasticity and overall performance is due to poor performance in poor environments, but if linear plasticity values are negative, this can also indicate reduced performance in improved environments. We observed a tradeoff between linear plasticity for yield as a percentage of the environment mean and average performance relative to the population in the 14 environments with the lowest mean yields out of the 31 environments that were used for plasticity analyses (Supplemental Figure 10C, Spearman *ρ* = –0.39). In contrast, we observed a positive asymptotic relationship between them in the 14 environments with the highest mean yields (Supplemental Figure 10D, Spearman *ρ* = 0.42). These relationships also held true when linear plasticity values were compared with mean yields across these environments (Supplemental Figure 10E, F). Plasticity values estimated in the 14 best environments were far more negative (–5.55 – 1.27; Supplemental Figure 10D, F) than plasticity values estimated in the 14 poorest environments (0.59 – 1.56; Supplemental Figure 10C, E). In all cases, mean parent release year was significantly positively associated with the measure of overall performance used (*p* < 1.0 × 10^−15^) when controlling for linear plasticity. This indicates that, relative to the population, newer hybrids are responding less negatively to poor environments and more positively to improved environments than older hybrids (Supplemental Figure 10).

The measures of linear plasticity we have presented above consider all factors that differ between the environments tested, but do not attribute the phenotypic variation to any specific environmental factor. However, in some cases it may be desirable to modulate the level of plasticity in response to a specific environmental factor, such as the level of nitrogen fertilization. On account of the significant input costs^22^, greenhouse gas production^23^, and negative environmental^24^ and human health impacts^25^ associated with the production and agricultural use of synthetic nitrogen fertilizers, there has recently been increased interest in decreasing their use, but not at the cost of decreasing yield. Developing maize hybrids with yields that remain stable under reduced levels of nitrogen fertilization via traditional selection methods requires accurate measurement of the nitrogen plasticity under environmental conditions relevant to the target environments^26^. Because the true values of phenotypes are often unknown, repeatable measurements are often used in place of accurate measurements. As the genetic repeatability of measurements for a trait increases, the effectiveness of selection also generally increases, given the same selection intensity^27^. We found the repeatability of nitrogen plasticity for yield to be quite low across location-years (− 0.07 ≤ *ρ* ≤ 0.15, Figure 2A, B) and even within a location-year (− 0.02 ≤ *ρ* ≤ 0.17, Supplemental Figure 2).

The low repeatability of nitrogen plasticity measurements in this study may be due to several causes. First, nitrogen plasticities were calculated using yield estimates from only two replicates of each genotype in each of the three nitrogen levels in a location-year, so the estimates of the genotype mean yields and resulting nitrogen plasticities may be noisy. However, two replicates in each environment per genotype were sufficient to repeatably measure complex traits such as yield and ear height in high nitrogen-fertilization environments with Spearman rank correlations within an environment of 0.30 ≤ *ρ* ≤ 0.77 and 0.20 ≤ *ρ* ≤ 0.70, respectively. In 33% of cases, nitrogen plasticities correlated better between location-years using two replicates per genotype in each environment than within a location-year using one replicate per genotype in each environment (Figure 2A, Supplemental Figure 2). This indicates that increasing the number of replicates can improve the repeatability of yield nitrogen plasticity by increasing the repeatability of the grain yield measurements. It is unclear how many replications of a genotype per environment would be required to measure nitrogen plasticity sufficiently repeatably to effectively select for nitrogen stability.

In this study, we showed that hybrid maize grain yield and linear plasticity have increased together during U.S. hybrid maize breeding between 1934 and 2000 (Figure 3B), and that performance has increased in all environments (Supplemental Figure 6). Improved performance in all environments has contributed to the increase in linear plasticity with overall yield (Supplemental Figure 4). There are two divergent paths toward increasing the environmental sustainability of maize production specifically and crop production generally. First, breeders can seek to reduce the required inputs given the same land area. This approach would require breeding lines with yields that are stable under reduced fertilizer and irrigation inputs. We found nitrogen plasticity to be poorly repeatable within and across location-years (Figure 2, Supplemental Figure 2), making selection for nitrogen-stable lines, a key component of this strategy, difficult. Though adding replicates or additional nitrogen fertilization rates may increase the repeatability of this trait, it would also increase the cost of field trials used for selection. However, because the performance in poor environments has not decreased despite increases in plasticity (Figure 3A–B, Supplemental Figure 6) and performance has improved across all environments (Supplemental Figures 5, 4), this may not be necessary. The second possible approach is to focus on breeding for increased production in areas of high potential yield. The observed improvements in performance across all environments suggests that selection for performance in high-yield-potential environments may also improve performance in marginal environments. The environmental impact per bushel of corn produced tends to be lower in high-productivity systems than in low-yield systems^28^. Ideally, this would also allow for better containment and mitigation of environmental impacts such as nitrate leaching via concentrated efforts. This approach would create incentives to breed for hybrids that not only can respond strongly to highly favorable environments, but also can tolerate high-intensity production practices such as high-density planting.

## Methods

### Maize field experiments and data processing

Of the 122 hybrid entries in this study, seven are hybrids which were commericially available at the time of the trials; 48 were drawn from a set of hybrids been repeatedly used in the multi-year, multi-site Genomes to Fields Initiative^29–32^; 36 are “Early Release” hybrids created using Corn Belt inbreds released prior to the 1950s, and the remainder are hybrids that have been previously utilized in other studies by our research group. The 115 non-commercial maize hybrids used in the Genomes to Fields Initiative trials, nitrogen yield trials, and “Early Release” hybrids were created by crossing 52 inbred maize lines released under a now expired Plant Variety Protection certificate, which we refer to as ex-PVP inbred lines, or released without intellectual property proection, typically by a public sector breeding program. Each inbred served as a parent for 1 to 13 hybrids. Individual hybrids are described in Supplemental Table 1. The mean parent release year for each hybrid was calculated as the mean of the release years of the parental inbred lines. The parental line release year was determined using the information available for the parental lines on the U.S. National Plant Germplasm System (NPGS) website (https://npgsweb.ars-grin.gov/gringlobal/search). When no release date or intellectual property right certificate was available, the earliest date of the receipt of the line or donation to NPGS was used. For detailed information on the parental inbred lines used to generate the hybrids in this study, please see Supplemental Table 2. Resequencing data for 47 of the 52 parental lines is available in Grzybowski et al.^33^, and the parental inbred lines used in this study can be matched to the inbred lines in the resequencing SNP set using Supplemental Table 3.

**Table 1.** Summary of irrigation provided and nitrogen fertilization rate(s) in each location-year.

| **Location-Year** | **Irrigation Provided (inches)** | **Nitrogen Fertilization Rates (lb N/acre)** | **Planting Date** | **Harvest Date** |
| --- | --- | --- | --- | --- |
| 2022 Scottsbluff | 16.9 (429 mm) | **Low:** 75 (84 kg N/ha)<br>**Medium:** 150 (168 kg N/ha)<br>**High:** 225 (252 kg N/ha) | May 19, 2022 | November 9 – 10, 2022 |
| 2022 North Platte No Irrigation (NI) | 0 (0 mm) | **Low:** 75 (84 kg N/ha)<br>**Medium:** 150 (168 kg N/ha)<br>**High:** 225 (252 kg N/ha) | May 18, 2022 | October 19 – 24, 2022 |
| 2022 North Platte Limited Irrigation (LI) | 4.3 (109 mm) | **Low:** 75 (84 kg N/ha)<br>**Medium:** 150 (168 kg N/ha)<br>**High:** 225 (252 kg N/ha) | May 17, 2022 | October 26 – 28, 2022 |
| 2022 North Platte Full Irrigation (FI) | 8.6 (218 mm) | **Low:** 75 (84 kg N/ha)<br>**Medium:** 150 (168 kg N/ha)<br>**High:** 225 (252 kg N/ha) | May 17, 2022 | October 31 – November 1, 2022 |
| 2022 Lincoln | 0 (0 mm) | **Low:** 75 (84 kg N/ha)<br>**Medium:** 150 (168 kg N/ha)<br>**High:** 225 (252 kg N/ha) | May 22, 2022 | October 10, 2022 |
| 2022 Missouri Valley | 0 (0 mm) | **Medium:** 175 (196 kg N/ha) | April 29, 2022 | October 11, 2022 |
| 2022 Ames | 0 (0 mm) | **Low:** 75 (84 kg N/ha)<br>**Medium:** 150 (168 kg N/ha)<br>**High:** 250 (280 kg N/ha) | May 22 – 23, 2022 | October 12 – 16, 2022 |
| 2022 Crawfordsville | 0 (0 mm) | **Low:** 75 (84 kg N/ha)<br>**Medium:** 150 (168 kg N/ha)<br>**High:** 225 (252 kg N/ha) | May 11, 2022 | October 7, 2022 |
| 2023 North Platte No Irrigation (NI) | 0 (0 mm) | **Medium:** 150 (168 kg N/ha) | May 10, 2023 | October 19 – 20, 2023 |
| 2023 North Platte Limited Irrigation (LI) | 4.5 (114 mm) | **Medium:** 150 (168 kg N/ha) | May 10, 2023 | October 19 – 20, 2023 |
| 2023 Lincoln | 0 (0 mm) | **Low:** 75 (84 kg N/ha)<br>**Medium:** 150 (168 kg N/ha)<br>**High:** 225 (252 kg N/ha) | May 16, 2023 | October 23, 2023 |
| 2023 Missouri Valley | 0 (0 mm) | **Medium:** 160 (179 kg N/ha) | May 10, 2023 | September 25, 2023 |
| 2023 Ames | 0 (0 mm) | **Low:** 75 (84 kg N/ha)<br>**Medium:** 150 (168 kg N/ha)<br>**High:** 225 (252 kg N/ha) | May 19, 2023 | October 19, 2023 |
| 2023 Crawfordsville | 0 (0 mm) | **Low:** 75 (84 kg N/ha)<br>**Medium:** 150 (168 kg N/ha)<br>**High:** 225 (252 kg N/ha) | May 4, 2023 | October 2, 2023 |

We defined a location-year as a unique combination of location, year, and irrigation level, as irrigation was mostly confounded with location and year in our study (Figure 1A). Table 1 describes the irrigation provided and the nitrogen fertilization rate(s) in each location-year. We defined an environment as a unique combination of location-year and nitrogen fertilization level. Within each environment, a randomized complete block design with two blocks was utilized. All field plots were four-row plots with 30” row spacing, with a targeted planting density of 36,852 – 39,710 plants/acre (91,063 – 98,125 plants/ha). In Scottsbluff during the 2022 growing season, all field plots were 22.5 feet (6.86 meters) long (planted length). In all other location-years, all field plots were 17.5 feet (5.33 meters) long. Figure 1A shows a geographic overview of the experimental design employed in this study. The map of precipitation in Nebraska and Iowa in Figure 1A was produced using data from gridMET^34^ and the packages AOI^35^, climateR^36^, zonal^37^, and sf^38^. We county mean daily precipitation over the period of November 1, 2021 – October 31, 2023 over which coincides with the approximate end of the growing season prior to this two-year study and the approximate end of the last growing season in this study. All other data visualizations were produced using the packages patchwork^39^, viridis^40^, scales^41^, and cowplot^42^. Code for all data visualizations is available in Supplemental Information 1.

In 2022, 84 maize hybrids were grown in six locations (Scottsbluff, Nebraska; North Platte, Nebraska; Lincoln, Nebraska; Missouri Valley, Iowa; Ames, Iowa; and Crawfordsville, Iowa). In 2023, 81 of these same hybrids and an additional 38 hybrids were grown in five locations (North Platte, Nebraska; Lincoln, Nebraska; Missouri Valley, Iowa; Ames, Iowa; and Crawfordsville, Iowa). Within a block, each genotype present during that growing season was represented at least once, with the exception of the environments in North Platte during the 2023 growing season because of field space constraints. Specifically, two genotypes were absent from the 2023 North Platte LI location-year and 36 hybrids were absent from the 2023 North Platte NI location-year. In North Platte, during the 2022 growing season, the non-irrigated trial was spatially separated from the irrigated trials. In Ames during the 2022 growing season, the 75 lb N/acre trial was spatially separated from the 150 and 250 lb N/acre trials.

For detailed information regarding agronomic practices, trait data collection, and thresholds for removing extreme values, please see Supplemental Information 2. Briefly, measurements included the number of standing plants in the center two rows of the plot, the height of attachment of the primary ear, and the height of the flag leaf of representative plants in each plot during the growing season. Additionally, the date that 50% of the plants in a plot reached anthesis and silking was recorded in the Scottsbluff, North Platte, and Lincoln locations. After the majority of plants in the field had senesced, four representative ears were harvested by hand from the center of the outer two rows of each plot, artificially dried, and phenotyped. Following hand harvest of the representative ears, the center two rows of the plot were mechanically harvested by a plot combine. The plot-level grain yield and flowering time data from the 2022 growing season were previously described in Shrestha et al^43^. Data from all environments were combined using R version 4.3.2^44^ using the packages tidyverse^45^, readxl^46^, lubridate^47^, and weathermetrics^48^. Supplemental Information 1 contains all code for combining the data and analyses described below. Ear phenotype values were averaged within each plot to produce a single value per plot. GDDs were estimated from temperature data collected by in-field weather stations (Watchdog 2700) and imputed with data from NASA POWER^49^ with *T*_*base*_ = 50°F and *T*_*opt*_ = 86°F. In the 2022 Crawfordsville location-year, the weather station malfunctioned, so the weather data from a Genomes to Fields weather station of the same brand and model located approximately 0.3 miles (0.5 km) away were used^50^. The yield per acre (15.5% moisture on a wet basis and 56-lb bushels) was calculated from the grain weight and grain moisture content estimated by the combine during the mechanical harvest of the center two rows of the plot. The distribution of values for each trait within a location-year was visually examined and extreme values were removed as described in Supplemental Information 2. In cases where a single extreme value for an ear phenotype caused an extreme mean value, the mean of the remaining values for the ear phenotype was used. Within each environment, all phenotype values were spatially corrected using SpATS^51^, fitting the plot identifier as the genotype random effect to obtain plot-level coefficients plus the model intercept used for all downstream analyses. Raw and spatially corrected values for all traits are available in Supplemental Table 4. Five hybrids were present in fewer than three environments, and were dropped for all downstream analyses, leaving 117 hybrids that were each observed in a minimum of four environments.

### Variance partitioning

The proportions of variance of each phenotype that could be attributed to environment, genotype, and the interaction between genotype and environment were estimated by fitting the mixed linear model described by Equation 1 where *Y*_*ijk*_ is the performance of the *i*th genotype in the *j*th environment, *g*_*i*_ is the random effect of the *i*th genotype, *e*_*j*_ is the random effect of the *j*th environment, *η*_*ij*_ is the random effect of the interaction between the *i*th genotype and the *j*th environment, and *ε*_*i jk*_ is the deviation of the *k*th individual of the *i*th genotype in the *j*th environment from the regression-fitted value. Model fitting was done using the lme4 R package^52^. The proportion of variance attributed to each effect was estimated by dividing the variance of the effect by the total phenotypic variance.

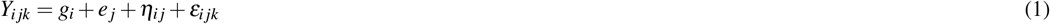

The proportion of variance in linear plasticity attributable to the general combining ability (GCA) of the parents was estimated as described above, substituting the model described by Equation 1 with Equation 2, where *B*_*ij*_ is the linear plasticity value for the hybrid with the *i*th ear parent and the *j*th pollen parent, *e*_*i*_ is the random effect of the *i*th ear parent, *p*_*j*_ is the random effect of the *j*th pollen parent, and *ε*_*ij*_ is the deviation of the linear plasticity value of the hybrid with the *i*th ear parent and *j*th pollen parent from the regression-fitted value.

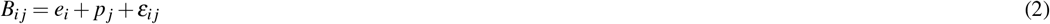

Because some parental lines were used as both ear parents and pollen parents in the creation of this hybrid population, we summed the proportions of variance attributed to *e*_*i*_ and *p*_*j*_ for a single estimate of the variance in linear plasticity attributable to the GCA of the parental inbred lines.

### Estimation of linear plasticity

The linear plasticity of each hybrid was estimated using the R package spFW^53^ ignoring spatial effects to fit the Finlay-Wilkinson joint regression model with the phenotype of interest as the response. Because the 2022 Lincoln location-year environments had significantly lower yield on a population level than all other location-years (Figure 1B), we did not include observations from the 2022 Lincoln environments in the data used to fit the model. The *b* value for each hybrid was extracted and regarded as the linear plasticity value. To make the data easier to interpret, 1 was added to all linear plasticity values.

The repeatability of linear plasticity values was estimated as the Spearman *ρ* between linear plasticity values for the hybrids obtained by fitting two Finlay-Wilkinson models, each using data from one block from each environment. The estimated Finlay-Wilkinson regression-fitted values for each hybrid were used to estimate whether there was a predicted rank order change between the hybrids when comparing the environment with the lowest population mean performance and the environment with the highest population mean performance. To estimate the relationship between overall mean trait value, linear plasticity, and mean parent release year, we performed an analysis of variance using lm() and anova() functions in R, using linear plasticity and mean parent release year as predictors and the overall mean trait value for the hybrid as the response. The overall rank of each hybrid across all studied environments was determined using a best linear unbiased prediction (BLUP) model with environment as a fixed effect. Overall rank was assigned in order of ascending BLUP values, with the hybrid having the minimum BLUP assigned a rank of 1.

For location-years where a positive relationship between grain yield and nitrogen fertilization rate was observed on a population level (2022 Scottsbluff, 2022 North Platte LI, and 2023 Ames), nitrogen linear plasticity was determined as described above using only data from the given location-year. A total of 31 hybrids were present only under medium nitrogen in the 2023 location-years, and were dropped from the analysis prior to estimating nitrogen plasticity. Correlation of nitrogen linear plasticity values between location-years was estimated using Spearman’s *ρ* with complete observations. The repeatability of nitrogen plasticity values was estimated as described above. The Spearman *ρ* between the linear plasticities estimated using each half of the dataset at a given location-year was estimated using all four location-years where a positive relationship between grain yield and nitrogen fertilization rate was observed on a population level.

For each hybrid, the hybrid mean yield within each environment as a percentage of the mean of all hybrids present in the environment was estimated. The line of best fit and associated 95% confidence intervals for this performance measure (as a function of environment) was estimated using the lm() function in R for each hybrid in the upper and lower 10% of all hybrids for either linear plasticity or yield across all environments. Linear plasticity for yield as a percentage of the environment mean was estimated using the lm() function in R in order to fit a model with the grain yield as a percentage of the environment mean as the response and the hybrid, the environment mean, and the interaction between the hybrid and the environment mean as predictors. The linear plasticity on a percent mean basis was taken as the environment mean interaction coefficient for the respective hybrid plus the population-level environment mean coefficient plus 1 to center the linear plasticity values near 1 for interpretability.

### Interaction importance scoring

The importance of the genotype-by-environment interaction between two hybrids in a pair of environments for informing selection decisions for a phenotype was scored using the following procedure. For each environment, an ANOVA was performed using the aov() function in R with genotype as the sole predictor. Tukey’s Honest Significance Difference (HSD) test was used to test all pairwise comparisons between hybrids. A pair of hybrids was considered to have a significant difference in this environment if the adjusted *p*-value was less than 0.05. For each pair of environments in which a hybrid pair was present, the pair of hybrids received a score of 0 if there was not a rank order change between the two environments. If the rank order of the two hybrids changed between the two environments, the pair of hybrids received a score equal to the number of environments in which there was a significant difference between the hybrids (0, 1, or 2). Figure 4B shows the performance and corresponding interaction importance scores for one hybrid pair in three pairs of environments. The overall score for the interaction between a pair of hybrids was calculated as the sum of the interaction importance scores across all environment pairs in which the pair of hybrids was present, and this score was normalized by dividing by the total possible score for the hybrid pair ([*n*_*E*_ × (*n*_*E*_ − 1)] where *n*_*E*_ is the number of environments in which a hybrid pair was present). Interaction scores for all hybrid pairs in all pairs of environments are available in Supplemental Table 5.

## Acknowledgements

The authors thank the members of the Iowa Crop Improvement Association, Jeff Golus, and Ramesh Kanna, and Deniz Istipiller for their efforts and assistance in conducting field trials and collecting ground truth data, and North Central Regional Plant Introduction Station (USDA-ARS and Iowa State University, Ames, Iowa, USA) for providing germplasm that facilitated this work. We are also grateful to Cole Hammett, Han Tran, Alliance Igiraneza, Isabel Sigmon, Daniella Tumisiime Norah, Chessie Ruester, Jordan Bares, Natasha Nguyen, Matthew Lowry, Vladimir Torres, Hector Palala, Kashish Syed, Halfen Khudada, Maddy Faber, Jennifer Lee, Esteban Heredia, Lauren McFadden, Timothy Moon, Peyton Stork, McKenna Murphy, Alexis Young, Allison Dagli, Charlie Vermeer, Elyse McElligott, Maggie Chapman, Haylee Thomas, Max Thomas, Janessa Allen, Hao Guo, Joseph Kaiser, Kyle Leonard, Grace Lowery, Mikaila Neuzil, Morgan Catron, Laura Boury, Korbyn Dewey, Nicholas Jenzano, Tyler Kennedy, Andrew Palar, Morgan Peak, Carson Ruen, Loryn Schaefer, Jon Swanson, Isaac Tharp, Ryan Wolf, and Joshua Woolfolk for their efforts in phenotyping ears. The authors are also grateful to Dr. Somak Dutta for his assistance in using the spFW() package to estimate linear plasticity, and to Réka Howard for her advice on the interaction importance scoring analysis.

## Author contributions statement

PSS and JCS conceived of the project. JMD, LMC, JT, LLC, KL, CU, and DKS conducted experiments and generated data. JMD, LMC, JT, KL, CU, and JCS processed data. JCS and PSS provided supervision, guidance, and feedback on the design of analyses and approaches. JMD conducted statistical analyses and visualized results. JMD drafted the paper with input from JCS. All authors contributed to the revision of the final manuscript.

## Funding

This research was supported in part by the U.S. Department of Agriculture, National Institute of Food and Agriculture under award numbers 2020-67021-31528, 2020-68013-30934, and 2023-70412-41087, and the AI Research Institutes program supported by NSF and USDA-NIFA under the AI Institute: for Resilient Agriculture, Award No. 2021-67021-35329. JMD is supported by the NSF Graduate Research Fellowship, Award ID 2034837.

## Conflict of interest statement

James C. Schnable has equity interests in Data2Bio, LLC and Dryland Genetics LLC and has performed paid work for Alphabet. Patrick S Schnable is a co-founder and CEO of Dryland Genetics, Inc and a co-founder and managing partner of Data2Bio, LLC. He serves as a member of the scientific advisory boards of Kemin Industries and Centro de Tecnologia Canavieira and is a recipient of research funding from Iowa Corn and Bayer Crop Science. He also serves as a consultant and/or expert witness in intellectual property disputes between seed companies. The authors declare no other conflicts of interest.

## Additional information

**Supplemental Table 1.**
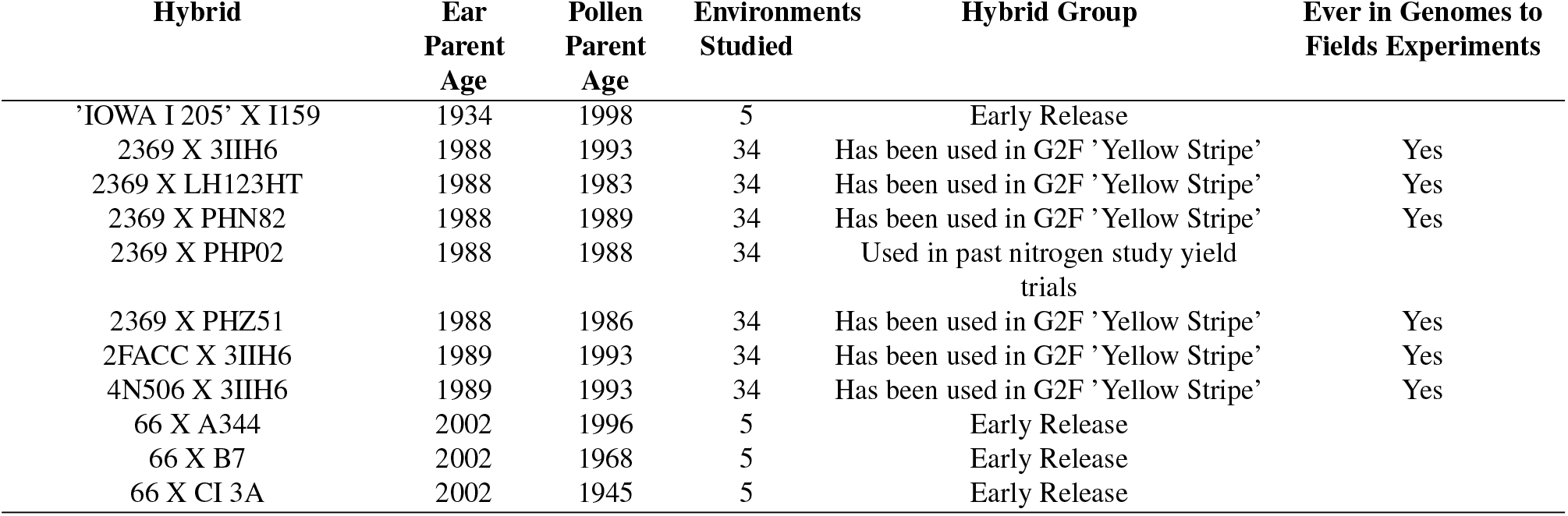
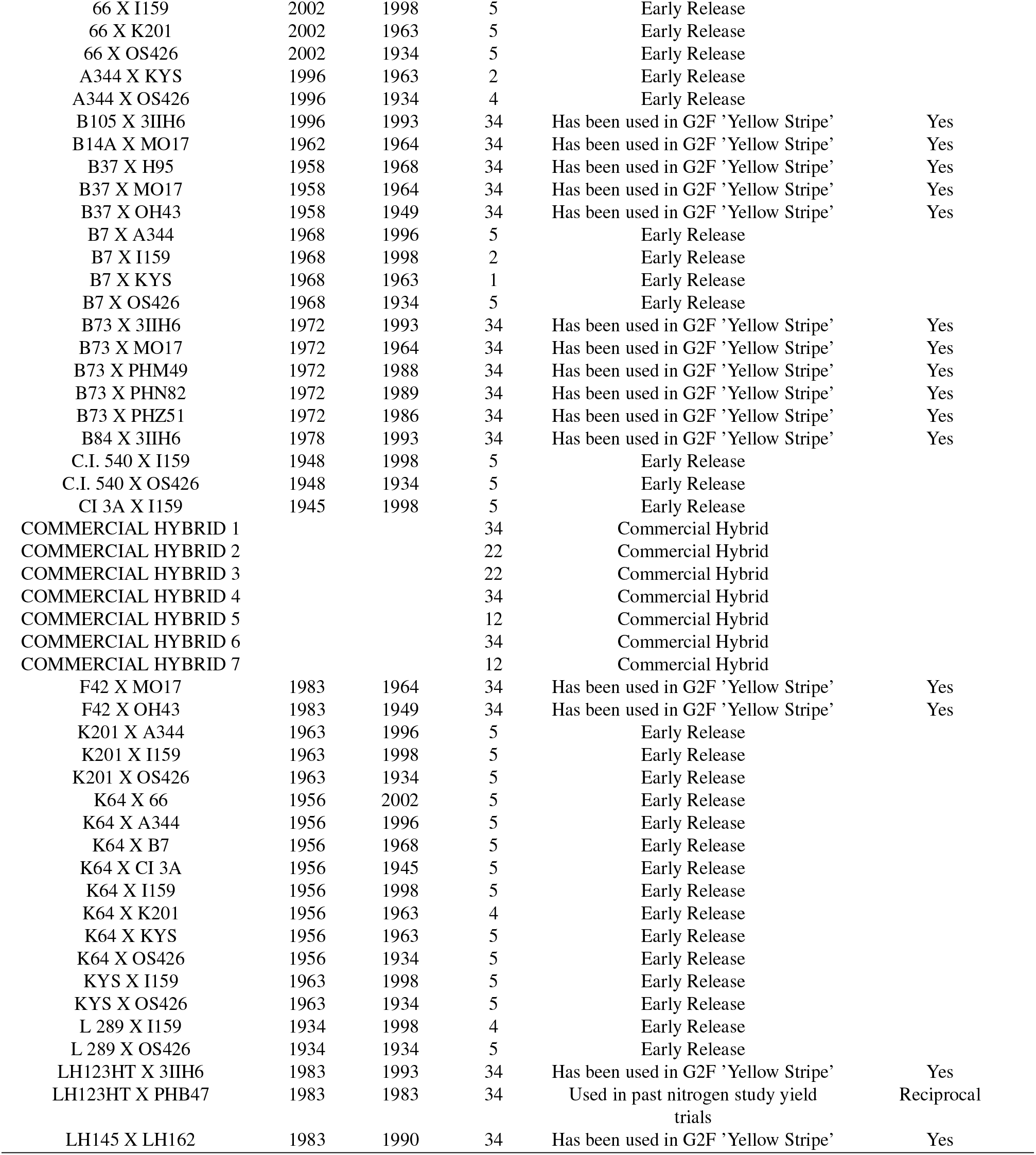
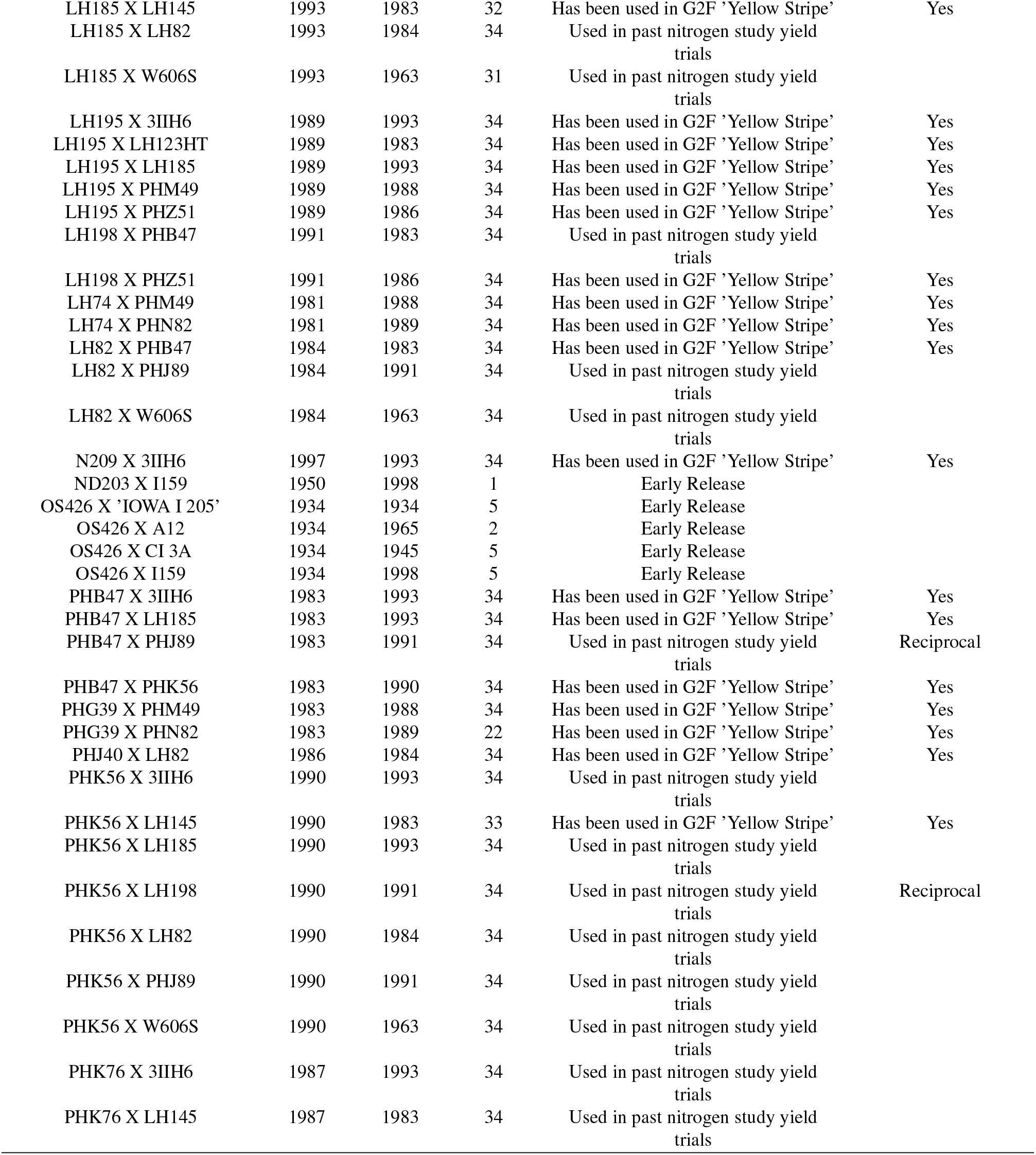
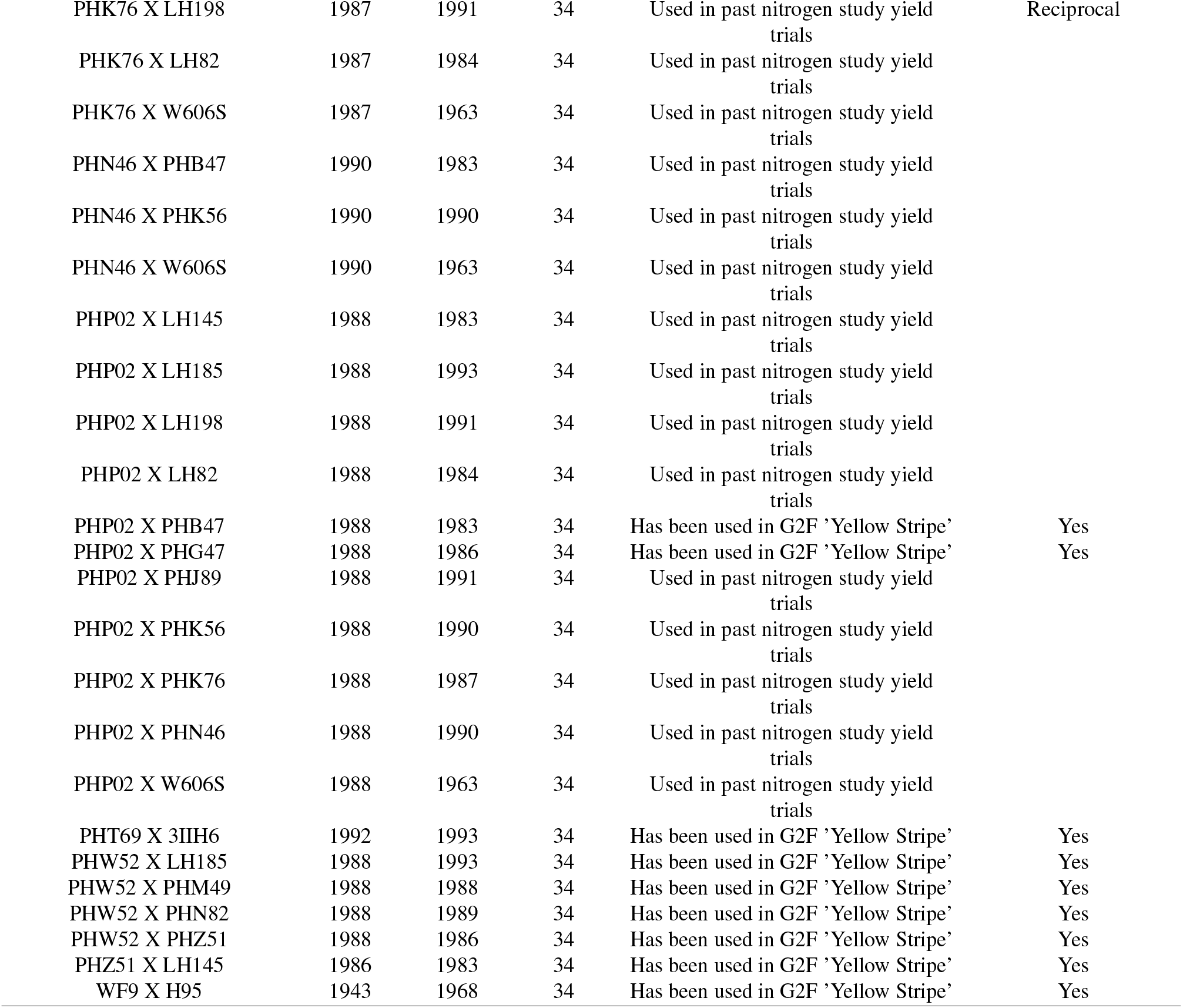
122 maize hybrids used in this study.

**Supplemental Table 2**. [Detailed information on parental lines used to create hybrid population used in this study](https://github.com/jdavis-132/hips/blob/master/outData/hybridParentInfo.csv).

**Supplemental Table 3**. [Conversion table linking names used for parental inbred lines in Supplemental Table 4 and the resequencing data available from Grzybowski et al. (2023)^33^](https://github.com/jdavis-132/hips/blob/master/outData/hybridParentalLinesConversionToGrzybowskiResequencing.csv).

**Supplemental Table 4**. [Raw and spatially corrected phenotype values](https://github.com/jdavis-132/hips/blob/master/analysis/HYBRIDS_2022_2023_SPATIALLYCORRECTED.csv).

**Supplemental Table 5**. [Interaction importance scores for all hybrid pairs in all pairs of environments for all phenotypes](https://github.com/jdavis-132/hips/blob/c3b2f240a4c6938a0033dd32d5f4c95a4d33d1ae/analysis/significantCrossovers.csv).

**Supplemental Information 1**. [Code used for data processing and analysis](https://github.com/jdavis-132/hips/blob/master/src/)].

**Supplemental Information 2**. [Detailed information on data collection and processing](https://github.com/jdavis-132/hips/blob/master/src/README_HYBRID_HIPS2022.html).

**Supplemental Figure 1.**
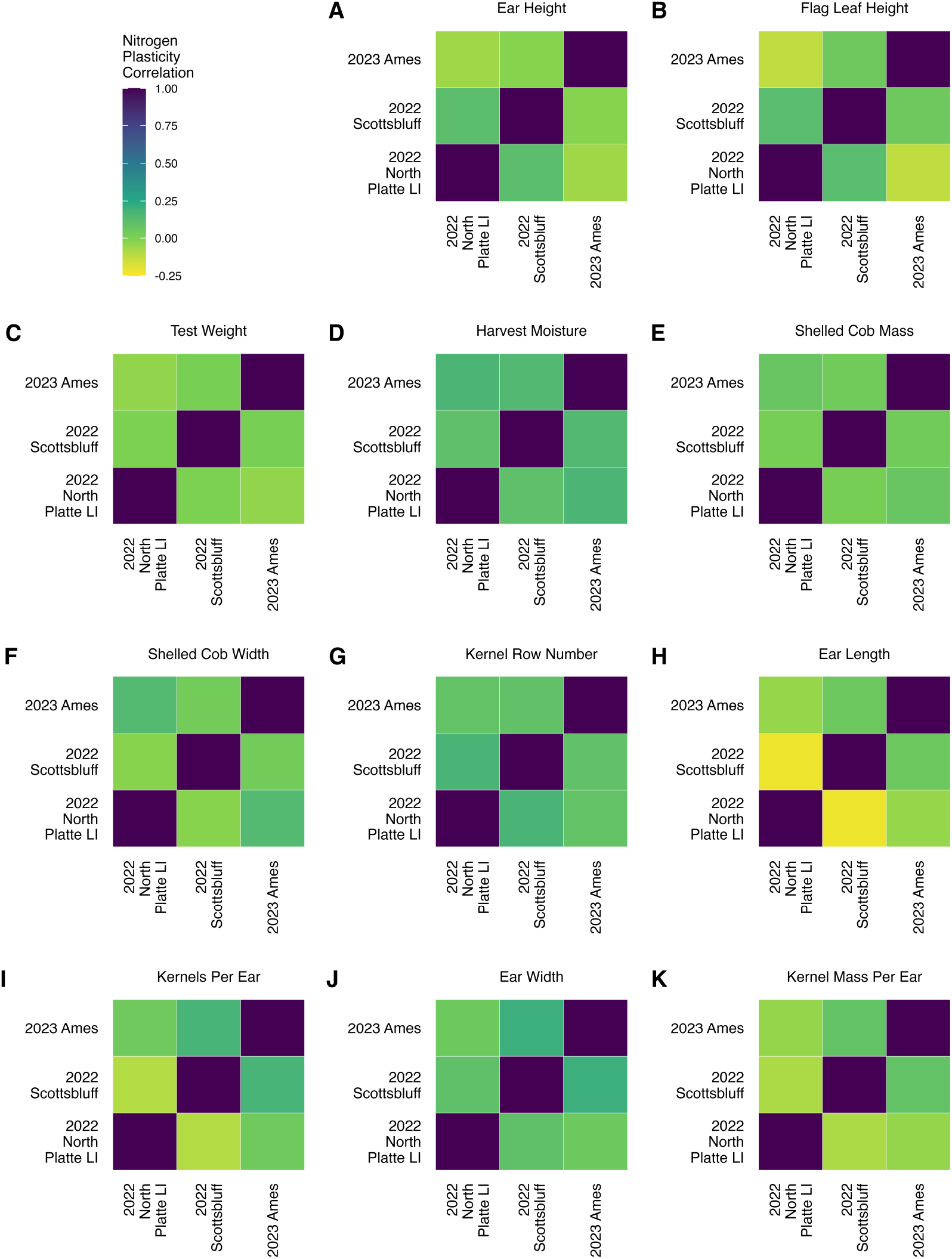
Nitrogen plasticity correlates poorly across location-years for all phenotypes collected in all location-years that were compared. **A – K)** Correlation of nitrogen plasticity for 11 phenotypes across three location years. Correlations were determined using the Spearman rank correlation of the linear nitrogen plasticity values estimated using data from all hybrids common to both location-years. Only location-years where a population-level positive relationship of yield with nitrogen fertilization was observed are shown.

**Supplemental Figure 2.**
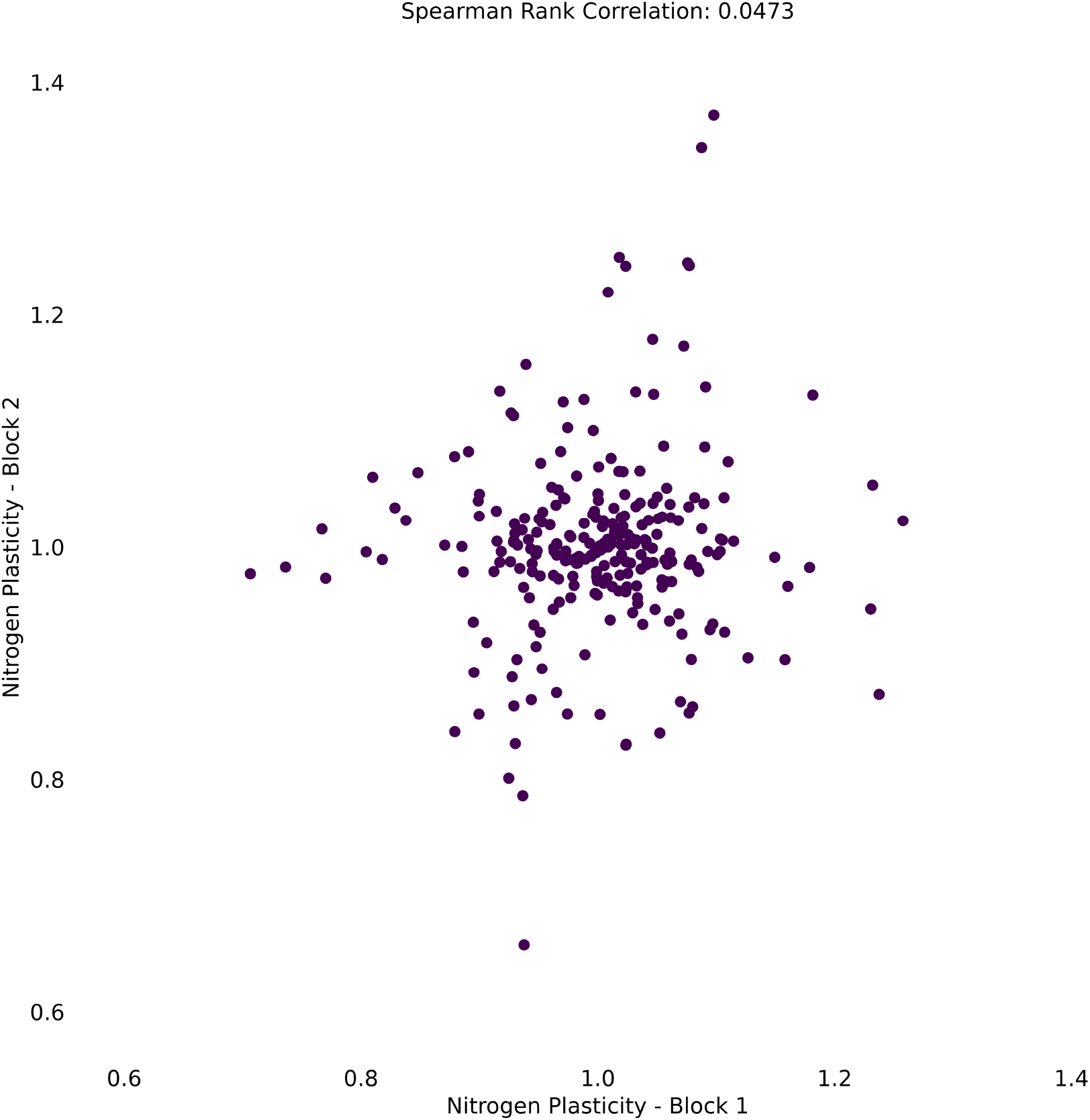
Nitrogen plasticity is poorly repeatable within a location-year. Nitrogen plasticity values were estimated for hybrids using 50% of the data in location-years in which a positive relationship with nitrogen fertilization rate was observed on a population level, compared to the equivalent estimates using the remaining 50% of the data. The data were split by block within an environment.

**Supplemental Figure 3.**
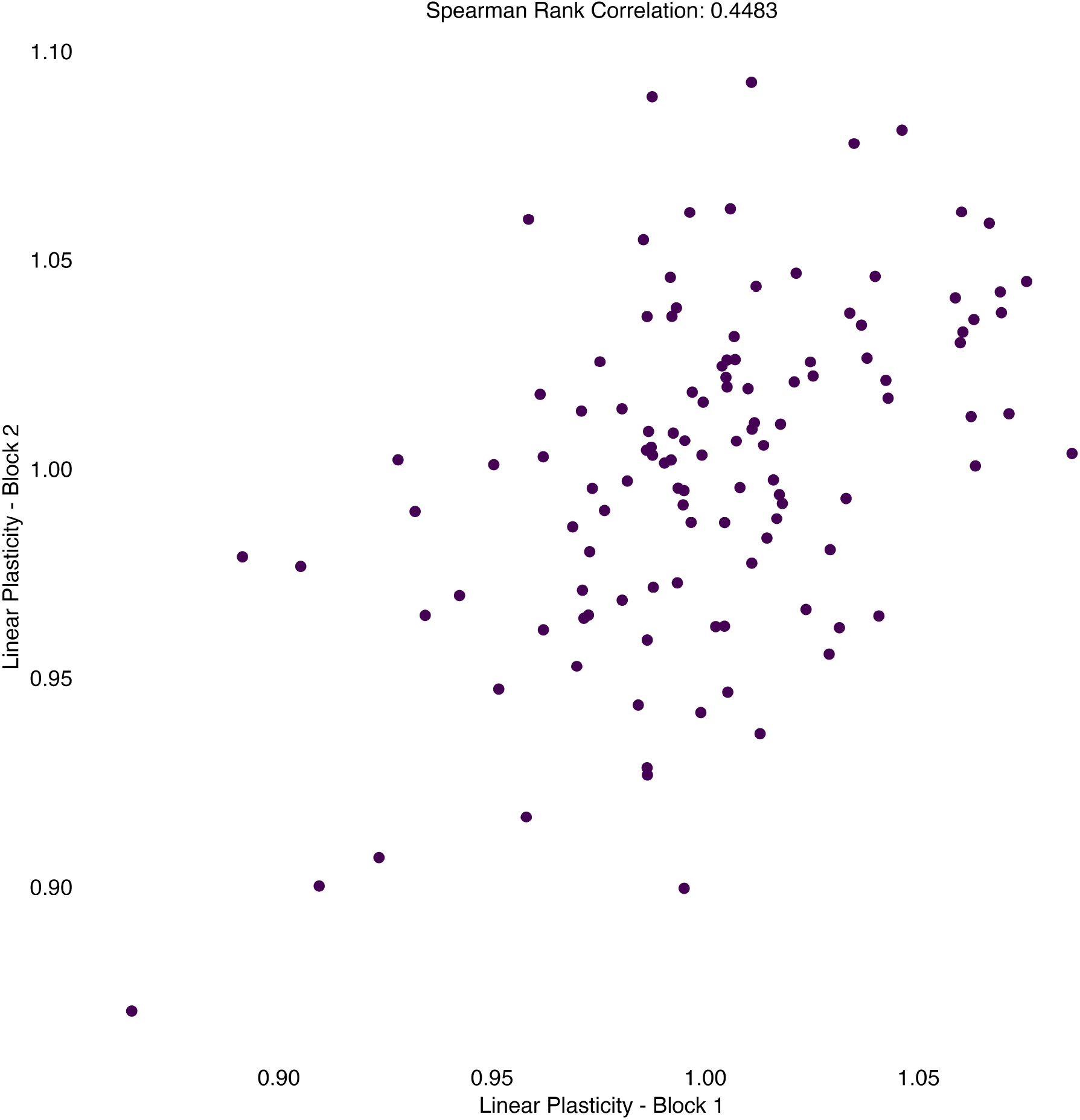
Linear plasticity values estimated for hybrids across all environments are moderately repeatable. The data were split by block within an environment.

**Supplemental Figure 4.**
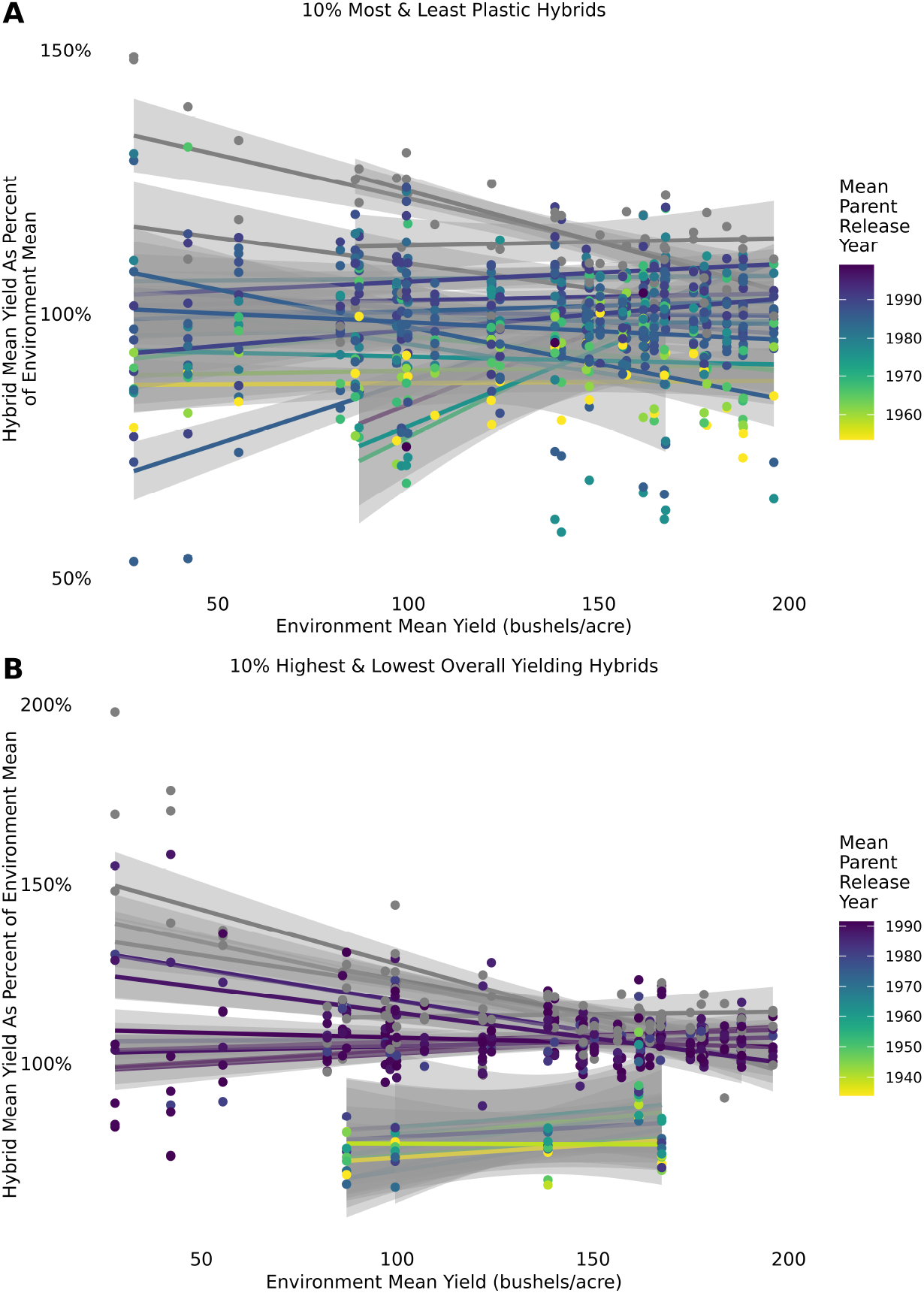
Performance of select hybrids as a percentage of the environment mean. **A)** The performance of the 10% most plastic and least plastic hybrids across all environments, with performance measured as the hybrid mean yield expressed as a percentage of the environment mean yield. Points represent hybrid means, and lines represent the line of best fit using the lm method and associated 95% confidence intervals. The color legend indicates mean parent release year. Gray indicates commercial hybrids for which both inbred parents are unknown. **B)** The performance of the hybrids with the highest and lowest 10% of yield BLUP values across all environments, with performance measured as the hybrid mean yield expressed as a percentage of the environment mean yield. Points represent hybrid means, and lines represent the line of best fit using the lm method and associated 95% confidence intervals. The color legend indicates mean parent release year. Gray indicates commercial hybrids for which both inbred parents are unknown.

**Supplemental Figure 5.**
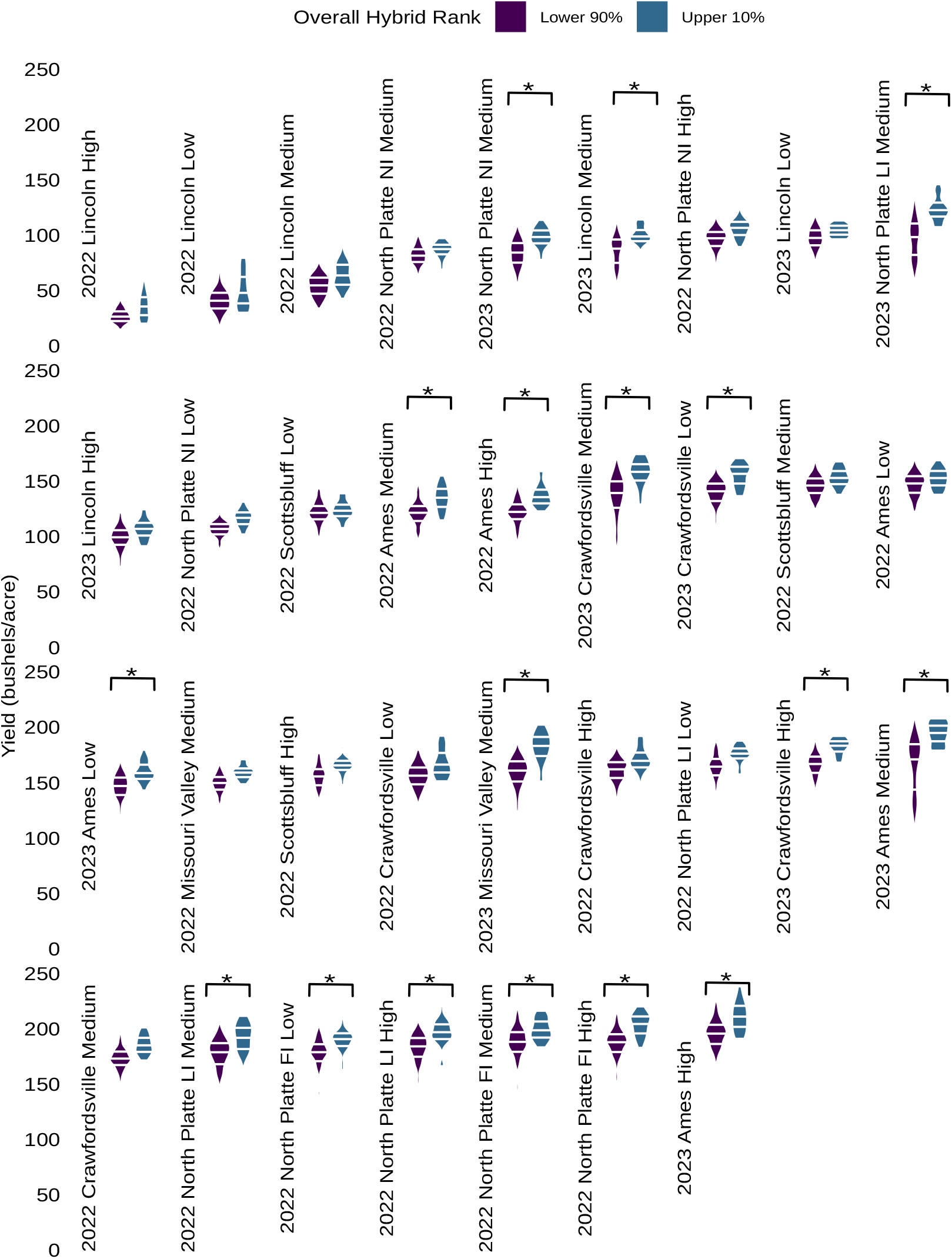
Yield of hybrids ranked in the upper 10% of hybrids for grain yield compared to the yield of all other hybrids by environment. Environments are shown in order of increasing environment mean yield from left to right and top to bottom. Asterisks denote environments where there was a significant difference (*p* < 0.05, Tukey’s HSD) between groups within the environment. Horizontal lines within each violin denote the 25^*th*^, 50^*th*^, and 75^*th*^ percentiles (Lower 90%: *n* = 106 – 147 plots; Upper 10%: *n* = 18 – 22 plots).

**Supplemental Figure 6.**
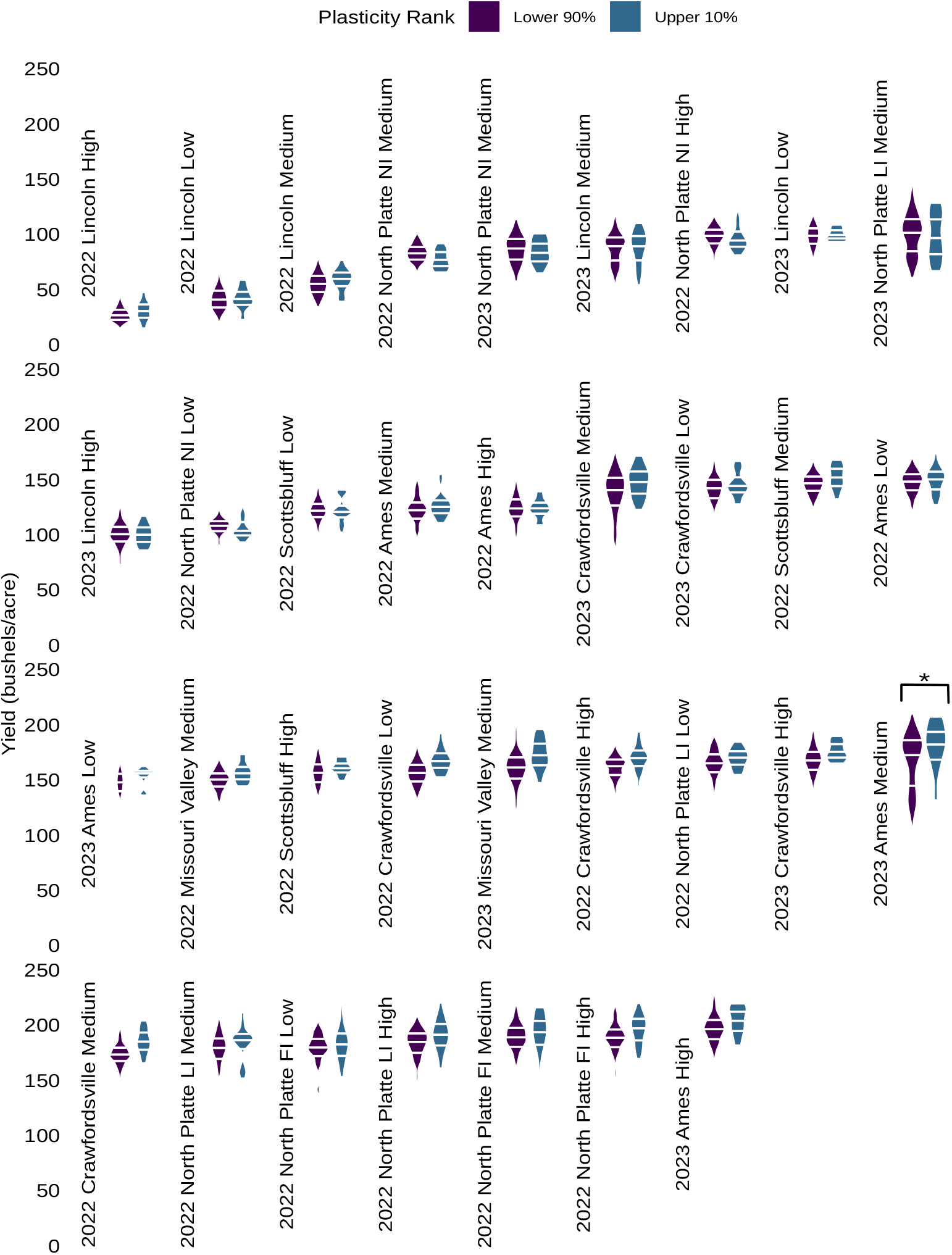
Yield of hybrids ranked in the upper 10% of hybrids for linear plasticity compared to the yield of all other hybrids by environment. Environments are shown in order of increasing environment mean yield from left to right and top to bottom. Asterisks denote environments where there was a significant difference (*p* < 0.05, Tukey’s HSD) between groups within the environment. Horizontal lines within each violin denote the 25^*th*^, 50^*th*^, and 75^*th*^ percentiles (Lower 90%: *n* = 114 – 206 plots; Upper 10%: *n* = 11 – 22 plots).

**Supplemental Figure 7.**
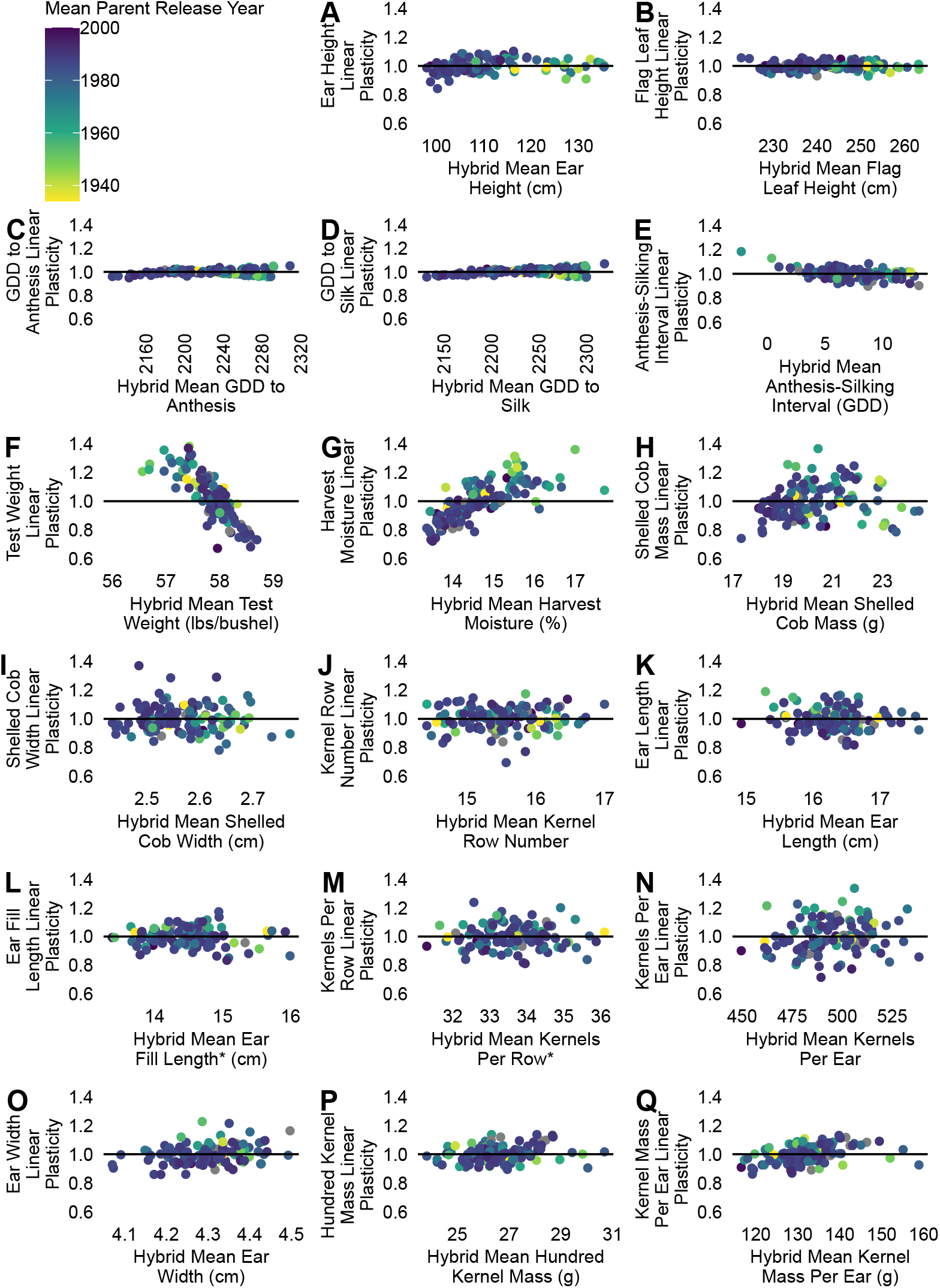
Relationship between Finlay-Wilkinson linear plasticity values across all environments, mean trait value, and mean parent release year is trait-dependent. **A – Q)** Finlay-Wilkinson linear plasticity values for each hybrid in this study versus their mean trait values across all environments, colored by their mean parent release year for 17 traits. Gray points represent commercial check hybrids for which the inbred parents are unknown. The black horizontal lines indicate a linear plasticity value of 1. GDD indicates growing degree days.

**Supplemental Figure 8.**
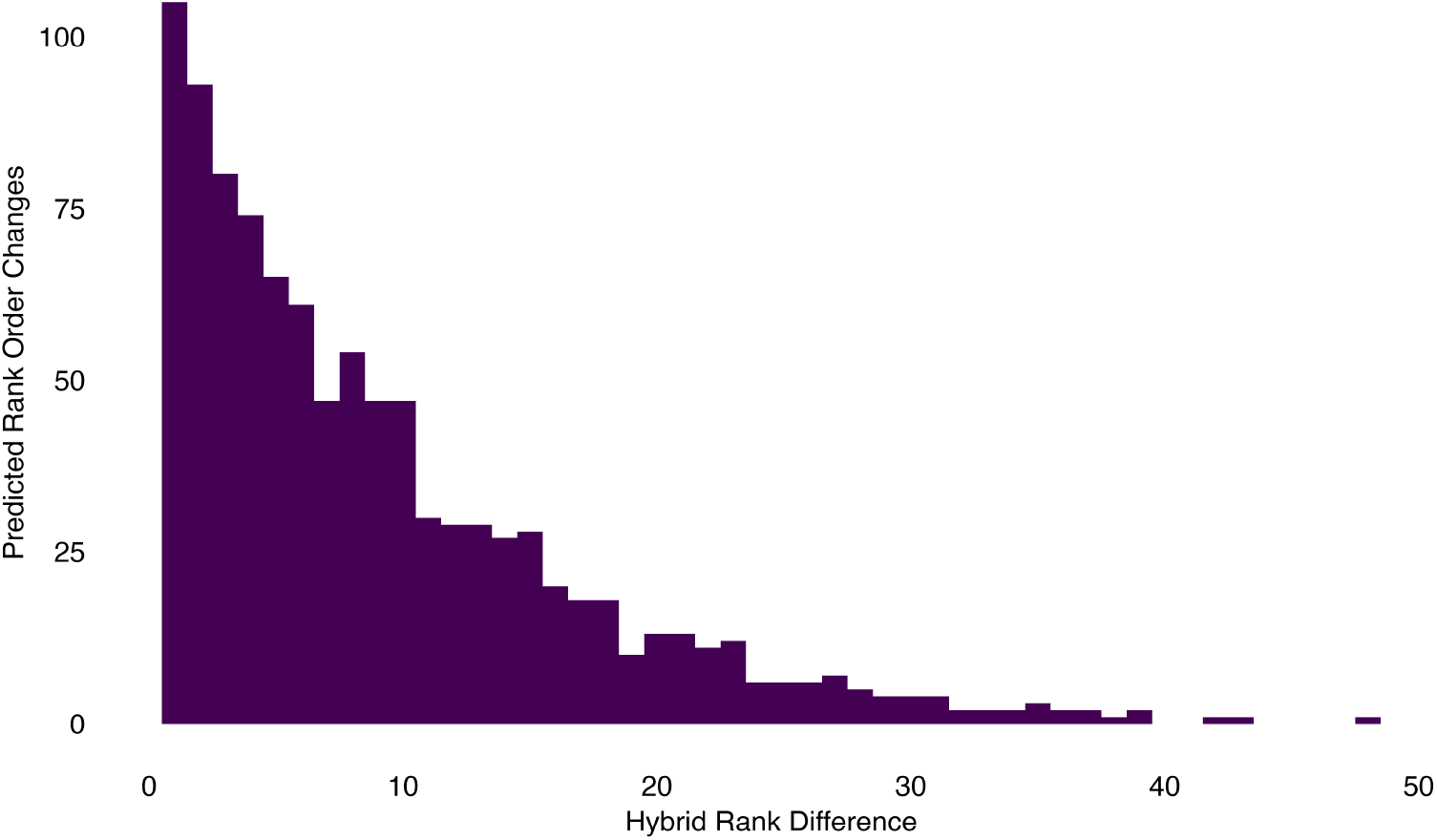
Distribution of the difference in overall hybrid ranks for the hybrid pairs where Finlay-Wilkinson regression predicted a rank order change in the population of hybrids studied.

**Supplemental Figure 9.**
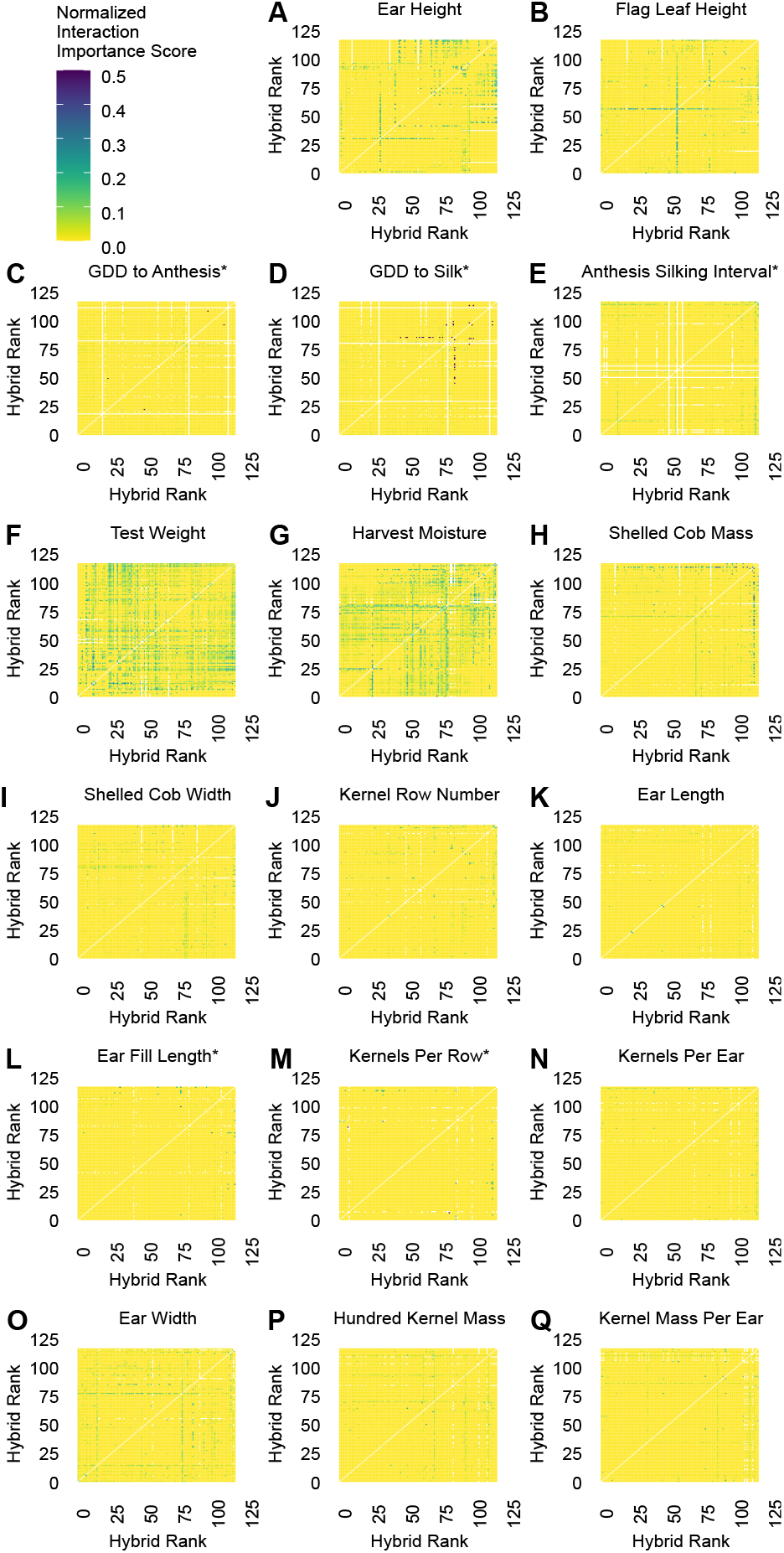
Pattern and level of interaction importance scores varied by trait. **A – Q)** Incidence matrices indicating the frequency with which a given pair of hybrids exhibited an interaction for the trait between two environments that represents a potentially important change in the selection decision between environments for 17 traits. For each hybrid and environment pair, an interaction received a score of 2 if the rank ordering of the hybrids changed between the two environments and there was a significant difference in yield between the hybrids in both environments. An interaction received a score of 1 if the rank ordering of the hybrids changed between the two environments and there was a significant difference in yield between the hybrids in one of the two environments. An interaction received a score of 0 otherwise. For each hybrid pair, interaction scores were summed across all environment pairs and divided by the total score possible for the hybrid pair based on the number of environments both hybrids were present in. Hybrids are ranked in order of ascending yield BLUP values fitting the environment as a fixed effect. Asterisks denote phenotypes measured only in a subset of environments. GDD indicates growing degree days.

**Supplemental Figure 10.**
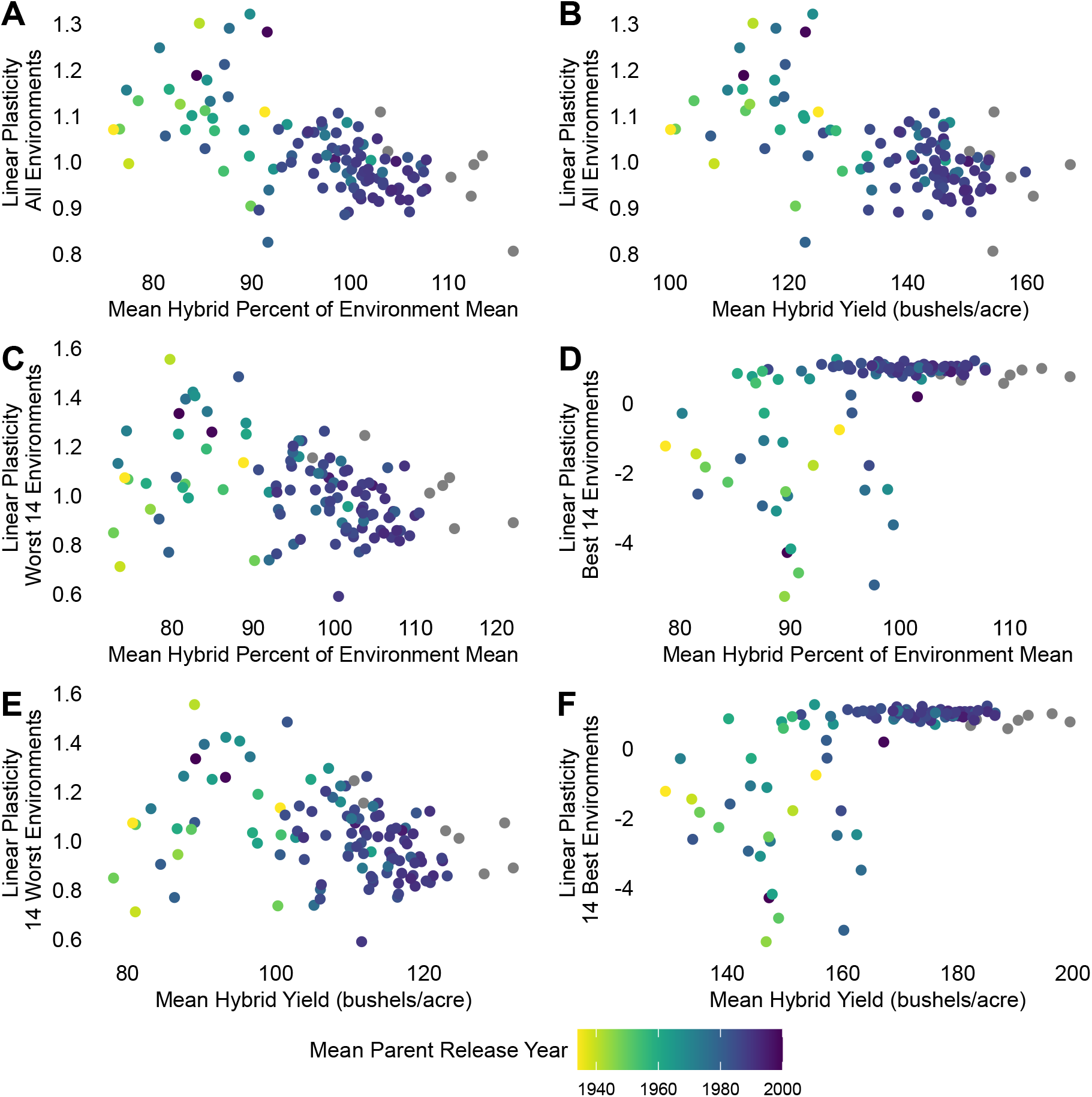
Linear plasticity for yield as a percentage of the environment mean yield shows a relationship with overall performance due to decreased yield response to improved environments. The color legend indicates mean parent release year. Gray points represent commercial check hybrids for which the inbred parents are unknown. **A)** Relationship between linear plasticity and average performance of hybrids relative to the population across all 31 environments used in plasticity analyses (Spearman *ρ* = -0.61). **B)** Relationship between linear plasticity and average yield of hybrids across all 31 environments used in plasticity analyses (Spearman *ρ* = -0.56). **C)** Relationship between linear plasticity and average performance of hybrids relative to the population in the 14 environments with the lowest environment mean yields used in plasticity analyses (Spearman *ρ* = -0.39). **D)** Relationship between linear plasticity and average performance of the hybrid relative to the population in 14 environments with the highest environment mean yields (Spearman *ρ* = 0.42). **E)** Relationship between linear plasticity and average yield of hybrids across the 14 environments with the lowest environment mean yields used in plasticity analyses (Spearman *ρ* = -0.36). **F)** Relationship between linear plasticity and average yield of hybrids across the 14 environments with the highest environment mean yields (Spearman *ρ* = 0.50).

## Notes

https://github.com/jdavis-132/hips

